# Identifying Modulators of the Post-Antibiotic Effect

**DOI:** 10.1101/2025.04.06.647494

**Authors:** Alexa L. Gilberti, Megan M. Tu, Kenneth Rachwalski, Madeeha I. Ali, Siobhan A. Cohen, Melike Akoglu, Alyssa C. Pollard-Kerning, Jinnette Tolentino Collado, Aya Sabbah, Franchesca Abou Said, Purnima Dela, Stephen G. Walker, John D. Haley, Eric D. Brown, Peter J. Tonge

**Affiliations:** Department of Chemistry, Stony Brook University, Stony Brook NY, 11794-3400, USA; Center for the Advanced Study of Drug Action, Stony Brook University, Stony Brook NY, 11794-3400, USA; Department of Biomedical Genetics, University of Rochester, Rochester, NY 14642, USA; Institute of Infectious Disease Research, McMaster University, Hamilton, ON, L8S 4L8, Canada; Biochemistry and Biomedical Sciences, McMaster University, Hamilton, ON, L8S 4L8, Canada; Chemistry Department, Farmingdale State College, Farmingdale, New York 11735, USA; Department of Oral Biology and Pathology, Stony Brook University, Stony Brook NY 11794-8702, USA; Department of Pathology, Stony Brook University, Stony Brook NY 11794-8691, USA

**Author notes:** To whom correspondence should be addressed: (EB), (PJT). A.L.G and M.M.T contributed equally to this work.

**Keywords:** post-antibiotic effect, LpxC, *Escherichia coli*, residence time, Keio collection

## Abstract

The post-antibiotic effect (PAE) is the delay in bacterial regrowth following antibiotic removal. It has important implications for dosing regimens since drugs that have extended activity following their elimination can be dosed less frequently, widening the therapeutic window. While the PAE has been associated with target vulnerability and the rate of target turnover, little is known about the genetic components that modulate the PAE. Here, we developed a high-throughput assay to screen the *Escherichia coli* Keio collection of ∼4000 deletion strains, identifying genes that enhance the PAE for CHIR-090, an inhibitor of UDP-3-*O*-(*R*-3-hydroxymyristoyl)-*N*-acetylglucosamine deacetylase (LpxC). This screen revealed approximately 400 gene knockouts that enhanced the PAE of CHIR-090. The list of PAE enhancers was enriched for genes involved in transmembrane transport and outer membrane synthesis. Notably, deletion of the *rfaE* gene, which is involved in lipopolysaccharide (LPS) biosynthesis, increased the PAE of the LpxC inhibitors CHIR-090 and LPC-058 by 2 h and 3 h, respectively. Consistent with this phenotype, co-treatment of wild-type *E. coli* with an RfaE inhibitor increased the PAE of CHIR-090 or LPC-058 by 1 h. To probe the mechanism of this interaction, we measured the rate of LpxC turnover and found that knocking out *rfaE* reduced its half-life by 2-fold, suggesting that disrupting RfaE increases the stability of LpxC, increasing target vulnerability and enhancing the PAE of LpxC inhibitors.

**SIGNIFICANCE:** Antibiotics are generally dosed at very high levels leading to unwanted side effects and non-compliance, which in turn results in the emergence of drug-resistant bacterial infections. The goal of this work was to develop strategies that will enable antibiotics to be dosed at lower levels, thereby improving safety and compliance. In the present work, we have screened 4,000 strains of *Escherichia coli* to identify compounds that result in an increase in the post-antibiotic effect (PAE), which is the delay in bacterial regrowth following antibiotic exposure and removal. Drugs that cause a PAE are dosed less frequently, and the method we describe will provide a new approach to developing safer drugs.

## INTRODUCTION

The post-antibiotic effect (PAE) is the persistent suppression of bacterial regrowth after the removal of an antimicrobial agent (1–3). Antibiotics that generate PAEs are dosed less frequently, improving patient adherence, reducing toxicity, and lowering treatment costs (2, 4, 5). For example, aminoglycosides, such as gentamicin (half-life ∼2 h), exhibit a significant PAE (2-7 h) against Gram-negative and Gram-positive bacteria, enabling extended-interval dosing strategies such as once-daily administration (6). In addition to aminoglycosides, fluoroquinolones, rifampin, and macrolides also generate extended PAEs, allowing less frequent dosing regimens without compromising therapeutic efficacy. In contrast, β-lactam antibiotics, such as cefazolin (half-life ∼2 h), have minimal or no PAEs against Gram-negative bacteria, necessitating frequent dosing such as every 6-8 h or continuous infusion to maintain sufficient drug concentrations (6). Notably, the extent to which a given antibiotic causes a PAE depends on the specific antibiotic-bacterium combination and growth conditions (6). Overall, the presence or absence of a PAE directly impacts dosing schedules and therapeutic strategies, underscoring its clinical importance.

Despite its therapeutic value, the mechanisms underlying PAEs remain poorly understood, limiting the development of approaches to design therapies with PAE in mind. Proposed mechanisms include reversible damage to cellular structures that requires recovery before bacterial growth can resume, antibiotic sequestration within bacterial cells, and the stability of the drug-target complex. Notably, a positive correlation between drug-target residence time and PAE has been observed for bacterial targets such as the ribosome (7), acetyl-CoA carboxylase (8), and UDP-3-*O*-(*R*-3-hydroxymyristoyl)-*N*-acetylglucosamine deacetylase (LpxC) in certain species (9, 10). Specifically, for *Pseudomonas aeruginosa* LpxC (paLpxC), prolonged drug-target binding is associated with an extended PAE (7, 9, 10). However, in *E. coli*, inhibitors of the corresponding enzyme (ecLpxC) produce little to no PAE due to rapid target turnover, reducing the vulnerability of ecLpxC (11). Thus, strategies to enhance the PAE by increasing target vulnerability or delaying cellular recovery pathways could improve antibiotic efficacy and reduce dosing frequency.

To investigate physiological factors that impact the PAE, we developed an innovative method to screen the Keio collection, containing 3,985 single-gene deletion mutants of nonessential genes in *E. coli* K-12 BW25113 (12), for mutations that prolong the PAE of antibiotics. Bacterial colonies were arrayed on filters and exposed to antibiotics by placing the filters onto drug-containing agar. Following a wash-out step, the colonies were pinned onto drug-free agar, and the time required for bacterial recovery was measured. Using CHIR-090, a compound that targets LpxC but does not induce a PAE in wild-type *E. coli* (11, 13), we identified 411 gene deletions that conferred a PAE on this compound, lengthening the recovery of bacterial growth compared to the no-drug control. The identified genetic enhancers of PAE were enriched for functions involved in cell motility, biofilm formation, and RNA metabolic processes. Notably, deletion of *rfaE*, which encodes an enzyme in the biosynthesis of LPS core precursor ADP-*L*-glycero-*D*-manno-heptose and is implicated in both cell motility and biofilm formation, doubled the half-life of LpxC protein and increased the PAE of CHIR-090 (14). Synergistic activity between the RfaE inhibitor **1** (compound-86 in Desroy *et al*. (15)) and the LpxC inhibitors CHIR-090 and LPC-058 (**Figure 1**) further validated the findings of our genetic screen. Overall, this study underscores the utility of this approach in uncovering novel targets to enhance the PAE of antibiotics.

**Figure 1:**
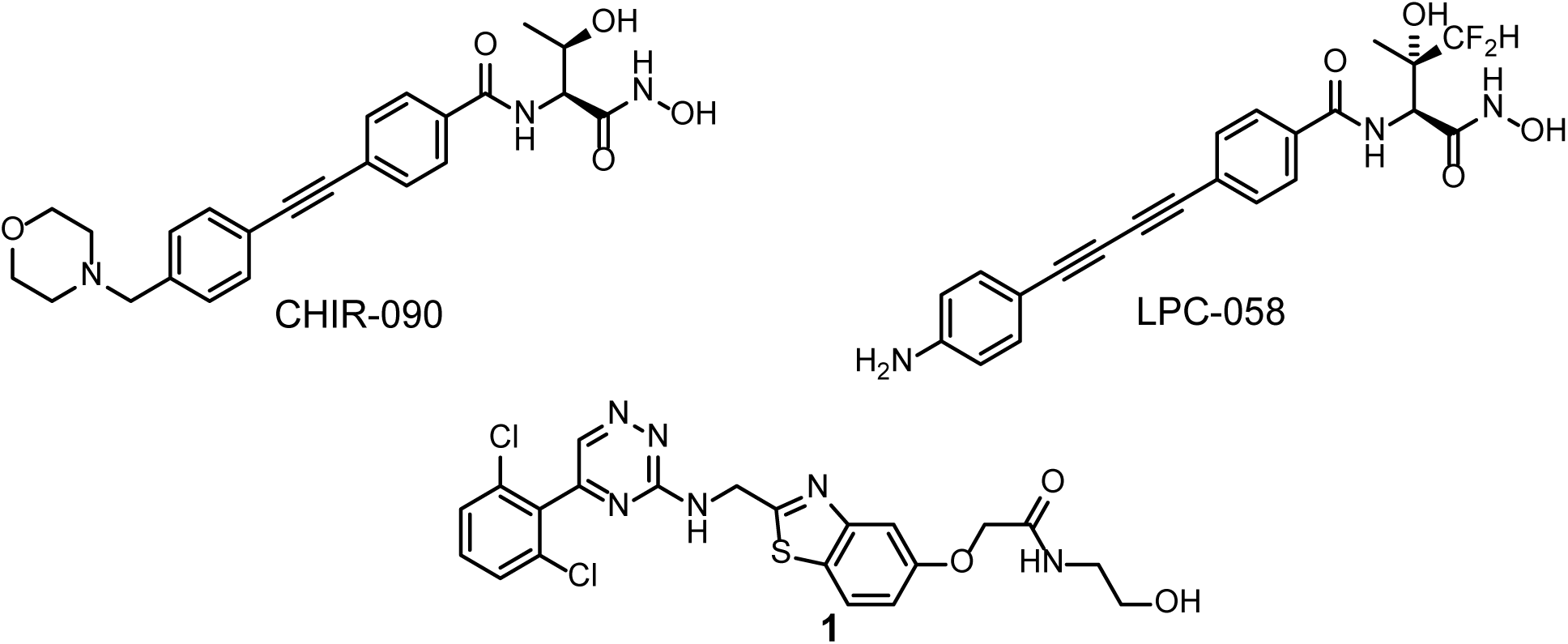
Compounds used in this work. Structures of the LpxC inhibitors CHIR-090 and LPC-058 and the RfaE inhibitor **1**.

## RESULTS

### Chemical-genetic screen of the Keio collection

To identify genetic determinants that influence the PAE of CHIR-090, we screened the Keio collection for mutants that enhanced the PAE. The Keio collection was arrayed onto nitrocellulose membranes in 384-density and grown overnight on LB-agar plates. The membranes were then transferred to LB agar plates supplemented with 16x MIC of CHIR-090 (MIC 0.29 µM) or no drug and incubated for 6 h (16). Following drug exposure, the colonies were manually pinned into 384-well plates containing PBS and subsequently pinned onto fresh LB agar plates. Colony growth was measured kinetically over 20 h using high-resolution scanners, and the lag time was calculated for each colony (**Figure 2A**).

**Figure 2:**
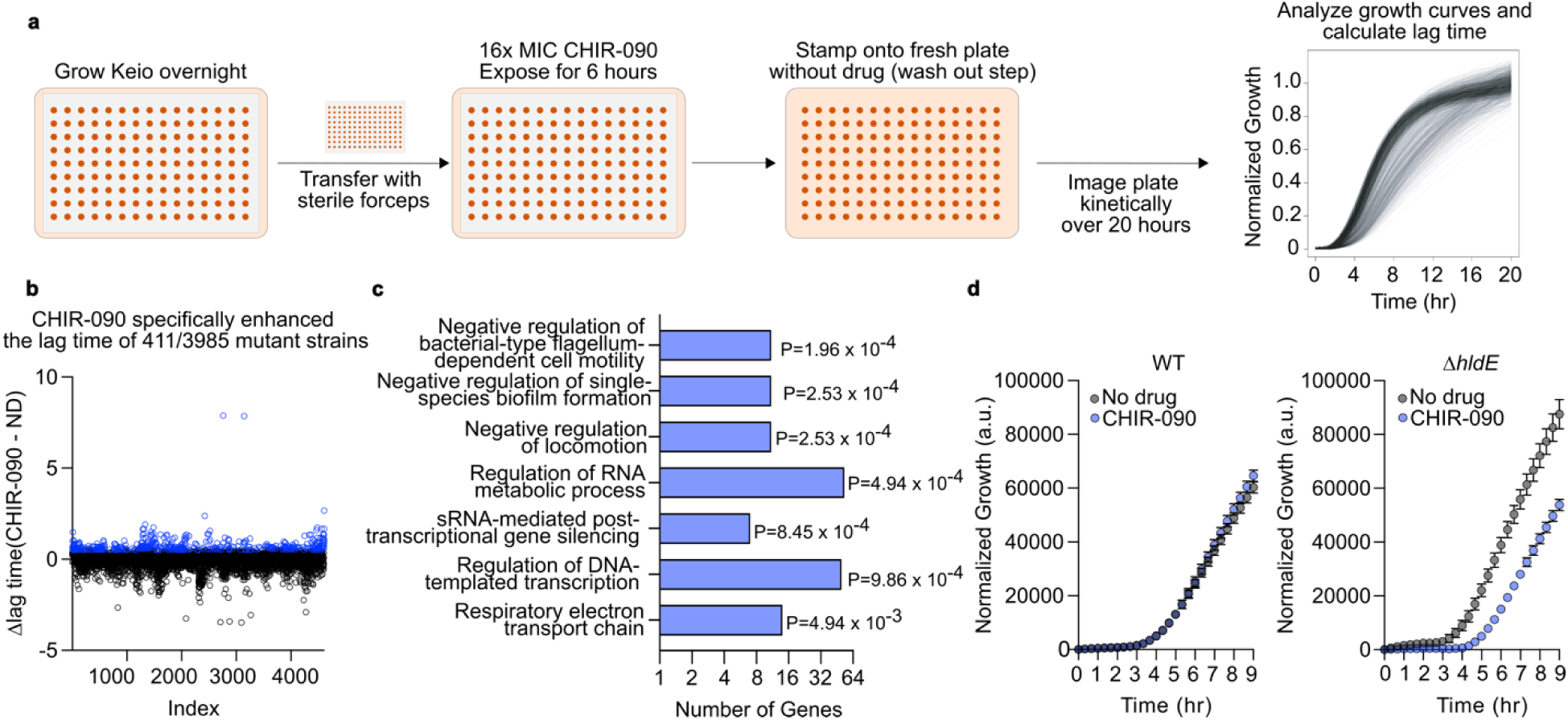
Enhancement of the lag time in growth of the Keio strains caused by CHIR-090. **(a)** Schematic for high-throughput PAE assay. The Keio collection of *E. coli* non-essential gene deletion strains is grown to endpoint in 384 colony density on PDMS filter paper. Mature colonies on filters are then transferred to an LB agar plate containing 16x the minimum inhibitory concentration of CHIR-090 or a no-drug control using sterile tweezers; colonies are exposed to the compound for 6 h. Subsequently, colonies are pinned onto fresh LB-agar without compound and plates are imaged using transmissive scans every 20 min for 20 h. Images are then be used to generate growth curves for each of the 384 mutants in the array, and growth curves are analyzed for enhancers of lag time. **(b)** Index plot depicting the change in lag time for each mutant after exposure to CHIR-090 relative to a no-drug control. Mutants whose lag time increased by 0.5 h (1 s.d. from the mean) specifically following CHIR-090 exposure are highlighted in blue. **(c)** Gene deletion mutants that had an increased lag time after CHIR-090 exposure were classified based on gene ontology (GO), and statistical enrichment was calculated using EcoCyc pathway-tools (28–30). The 7 most statistically enriched, non-redundant GO classifications are shown. Fisher’s exact test was used to calculate the *P*-value. **(d)** Growth curves of the mean growth of all colonies (WT) or Δ*rfaE* mutants after exposure to 16x MIC of CHIR-090 or the no-drug control.

The mean lag time of the Keio collection following exposure to CHIR-090 or no-drug was 3.38±0.17 h and 3.23±0.07 h, respectively. While not a strict measure of PAE, increased lag time is assumed to provide a reasonable proxy for the PAE. Therefore, the change in lag time in the presence of a drug relative to the no-drug control was determined for each mutant (**Figure S1**) (16, 17). A representative selection of strains that had lag times > 1 h are shown in **Table S1** and include lactaldehyde dehydrogenase *aldA* (18, 19), flagellar basal body rod protein *flgG* (20, 21), the drug transporter *yidY* (*mdtL*) (22), and heptose 7-phosphate kinase/heptose 1-phosphate adenyltransferase *rfaE* (14, 23–27). In total, 411 gene deletions experienced at least a 30 min increase in lag time after exposure to CHIR-090 (**Figure 2B**), and a subsequent analysis revealed several processes that may play a role in influencing the PAE. Specifically, deleting genes involved in cell motility, biofilm formation, locomotion, and RNA metabolic processes enhanced the PAE of CHIR-090 (**Figure 2C**). While the mechanistic connections between cell motility, biofilm formation, and the PAE are not immediately apparent, the relationship between RNA metabolism and the PAE is more evident. RNA processes are central to bacterial growth and recovery, and disruptions in RNA metabolism delay the time required for bacterial regrowth following antibiotic removal, thereby extending the PAE.

### Post-Antibiotic Effect of the *rfaE* Knockout Mutant in Liquid Media

Many of the genes identified in our screen, including those involved in cell motility and biofilm formation, are linked to cell envelope functions. We prioritized Δ*rfaE* for follow-up experiments due to its functional connection to LpxC through the biosynthesis of lipopolysaccharide (LPS), which is critical for maintaining outer membrane integrity and hence, impacts flagellar function and surface adhesion. Furthermore, inhibitors of RfaE have been reported, providing an opportunity to compare genetic and chemical perturbation on PAE generation. PAE measurements were subsequently conducted in liquid media using CHIR-090, which has a short residence time on LpxC (30 min), and the LpxC inhibitor LPC-058, which has a reported residence time of 12 h (**Figure 3**) (31), to evaluate the effect of drug-target residence time on PAE.

**Figure 3:**
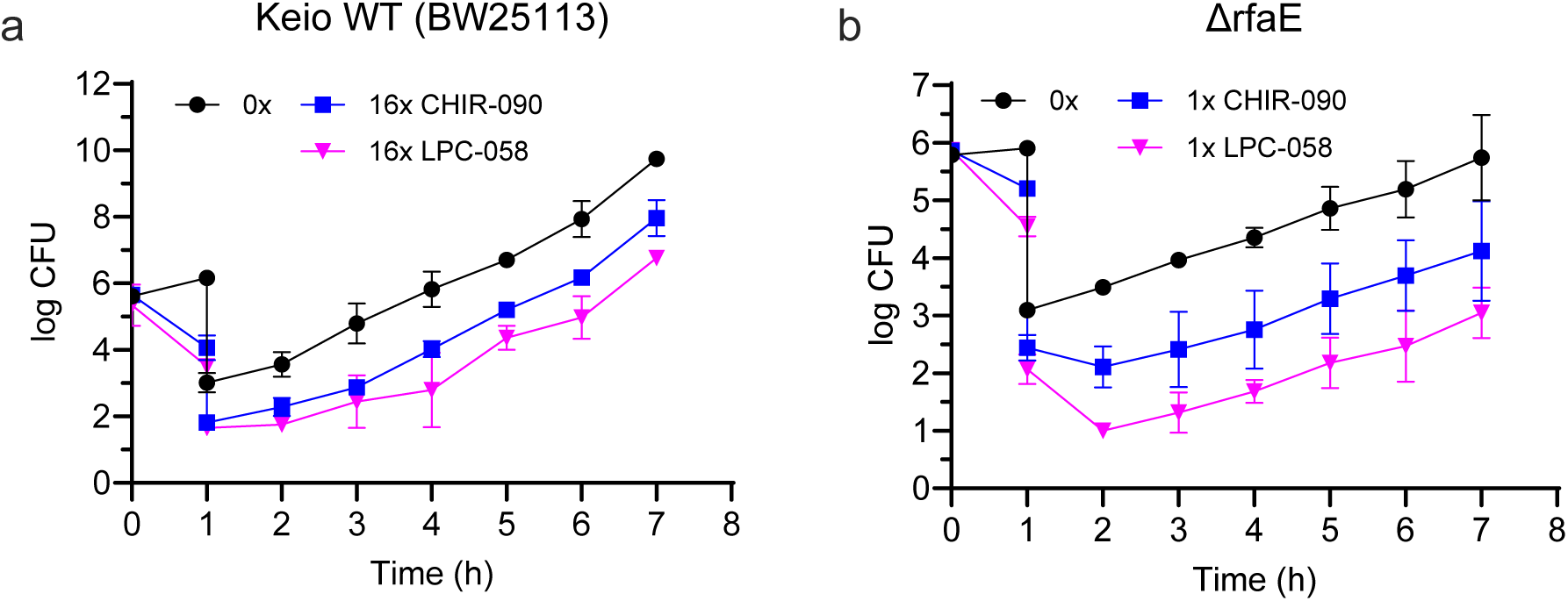
Post-antibiotic effect of CHIR-090 and LPC-058 on wild-type and Δ*rfaE* strains: Cultures of bacteria (10^6^ CFU/mL) were exposed to 1x, 2x, or 16x MIC of the compound for 30 min to 1 h followed by a 1:1000 dilution into fresh cation-adjusted Mueller-Hilton (CaMH) media at 37°C. Samples (100 μL) of the diluted cultures were then plated on Muller-Hinton agar plates every hour, and CFUs were enumerated following incubation of the plates at 37 °C for 16 h. (a) PAE of wild-type BW25113 induced by 16x MIC CHIR-090 and 16x LPC-058 after 1 h exposure (n=2). (b) PAE of Δ*rfaE* induced by 1x MIC CHIR-090 and 1x LPC-058 following a 45 min exposure (n=3). Experimental data points are the mean values of each experiment, and the error bars represent the standard deviation from the mean values.

The MIC values of CHIR-090 and LPC-058 against the *rfaE* mutant in liquid media were 0.04 μM and 0.06 μM, respectively, which are comparable to those obtained on solid media (**Table 1**, **Table S1**). However, although the wild-type strain could be exposed to 16x MIC of either CHIR-090 (MIC 0.16 μM) or LPC-058 (MIC 0.125 μM), time-kill experiments demonstrated that the maximum compound concentration for PAE experiments with the *rfaE* mutant was 1x MIC CHIR-090 or 1x MIC LPC-058 with an exposure time of 45 min (**Figure S2**). We then assessed the PAEs by monitoring the regrowth of bacteria after compound washout compared to the wild-type *E. coli*. Strains were exposed to either 1x or 16x MIC of the compounds for 45 to 60 min, after which the cultures were diluted 1:1000 fold into fresh media. Bacterial regrowth was monitored by enumerating colony-forming units (CFUs) over time, and the PAE was calculated by determining the time necessary for a 1 log increase in CFU post-washout compared to a non-treated culture with a vehicle of DMSO or nuclease-free water (7, 11). While no significant delay in bacterial regrowth was observed when the wild-type strain was treated with CHIR-090, LPC-058 generated a PAE of 1.3±0.7 h (**Table 1**). In addition, deleting *rfaE* increased the PAEs of CHIR-090 and LPC-058 to 2.1±0.8 h and 4.0±0.7 h, respectively (**Figure 3**).

**Table 1.** Microbiological data for the *E. coli* K12 Δ*acrAB*, Δ*tolC* pump mutant strain.

| **Table 1. Microbiological data for the *E. coli* K12 Δ*acrAB*, Δ*tolC* pump mutant strain** |  |  |  |  |  |  |  |  |
| --- | --- | --- | --- | --- | --- | --- | --- | --- |
| Strain | CHIR-090 | | CHIR-090 + 25 μM RfaE Inhibitor | | LPC-058 | | LPC-058 + 25 μM RfaE Inhibitor | |
| | MIC (μM)<sup>b</sup> | PAE (h)<sup>c</sup><br>16x MIC | MIC (μM)<sup>b</sup> | PAE (h)<sup>c</sup><br>16x MIC | MIC (μM)<sup>b</sup> | PAE (h)<sup>c</sup><br>16x MIC | MIC (μM)<sup>b</sup> | PAE (h)<sup>c</sup><br>16x MIC |
| WT<sup>a</sup> | 0.16 | 0 | n.d. | n.d. | 0.125 | 1.3±0.7 | n.d. | n.d. |
| Δ*rfaE* | 0.04 | 2.1±0.8<sup>d</sup> | n.d. | n.d. | 0.06 | 4.0±0.7<sup>d</sup> | n.d. | n.d. |
| Δ*acrAB*,<br>Δ*tolC* | 0.02 | 0 | 0.005 | 0.9±0.2 | 0.04 | 0.7±0.1 | 0.005 | 1.1±0.3 |

<sup>a</sup>WT, wild-type, *E. coli* K-12, BW25113. <sup>b</sup>MIC values were determined by the micro broth dilution method. Experiments were performed in triplicate, and the reported values are the average of the three independent experiments. <sup>c</sup>The PAE was calculated using a standard procedure, where the time required by the bacteria to recover 1 log CFU after washing out the inhibitor was compared to the culture (DMSO) (7, 11). n.d. not determined, as data was not collected and thus results not determined. <sup>d</sup>The PAE for $\Delta$ *rfaE* was determined at 1x MIC of CHIR-090 or LPC-058 after 45 min of exposure.

### Microbiological Experiments with an Inhibitor of RfaE

Next, we sought to phenocopy the genetic inactivation of *rfaE* through chemical inhibition. Desroy *et al*. described the development of **1** (**Figure 1**), an inhibitor of the RfaE kinase domain that further sensitized an efflux-deficient *E. coli* K12 Δ*acrAB*, Δ*tolC* strain to antibiotics such as erythromycin by inhibiting the biosynthesis of LPS (15). We thus assessed the impact of **1** on the microbiological activity of CHIR-090 and LPC-058 in the efflux-deficient strain. First, we confirmed the ability of **1** to enhance erythromycin sensitivity in the *E. coli* mutant Δ*acrAB*, Δ*tolC*, strain, as previously demonstrated. Indeed, the combination of **1** (12.5 μM) with a sub-MIC concentration of erythromycin (1.56 μM) was bactericidal as the RfaE inhibitor increased the sensitivity of the strain to erythromycin. We then examined how the sensitivity of *E. coli* Δ*acrAB*, Δ*tolC* to CHIR-090 or LPC-058 changed in the presence of **1**. When co-treated with **1** (25 μM), the MIC of CHIR-090 decreased 4-fold from 0.02 μM to 0.005 μM, and the MIC of LPC-058 decreased 8-fold from 0.04 μM to 0.005 μM.

We then determined the PAEs of CHIR-090 and LPC-058 in the *E. coli* Δ*-acrAB*, Δ*tolC* strain. In the absence of **1**, the PAE for 16x MIC CHIR-090 was 0 h, which increased to 0.9±0.2 h in the presence of 25 μM **1** (**Table 1**, **Figure 4**). Similarly, the PAE for 16x MIC LPC-058 in *E.coli* Δ*-acrAB*, Δ*tolC* was 0.7±0.1 h, which increased to 1.1±0.3 h in the presence of 25 μM **1** (**Table 1**, **Figure 4**). Although these experiments were performed in an efflux pump mutant strain, the findings demonstrate that chemical inhibition of RfaE using **1** can phenocopy the effects of *rfaE* genetic disruption, enhancing the PAE of the LpxC inhibitors.

**Figure 4:**
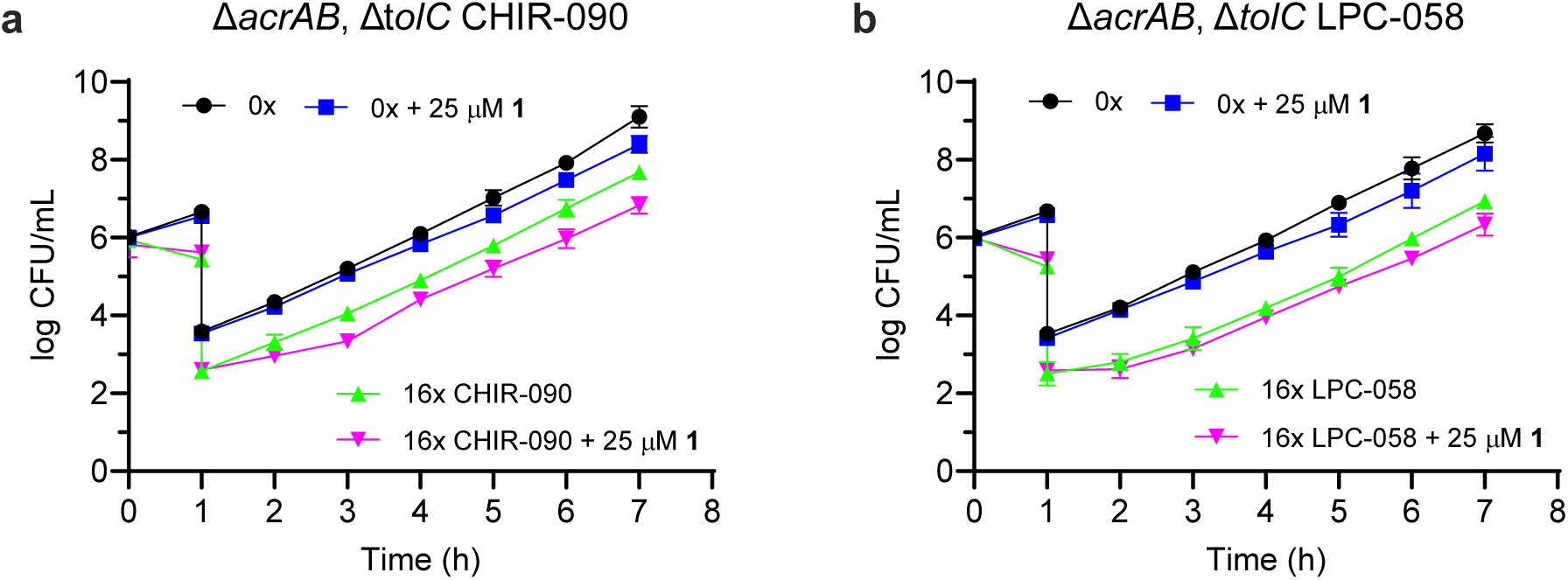
Post-antibiotic effect of CHIR-090 and LPC-058 on Δ*acrAB*, Δ*tolC* with compound 1. Cultures of bacteria (10^6^ CFU/mL) were exposed to 16x MIC of compound for 1 h followed by a 1:1000 dilution into fresh cation-adjusted Mueller-Hilton (CaMH) media at 37°C. Samples (100 μL) of the diluted cultures were then plated on Muller-Hinton agar plates every hour, and CFUs were enumerated following incubation of the plates at 37°C for 16 h. (a) PAE of *ΔacrAB, ΔtolC* induced by 16x MIC CHIR-090 following a 1 h exposure (n=3). (b) PAE induced by LPC-058. Experimental data points are the mean values of each experiment, and the error bars represent the standard deviation from the mean values.

### LpxC Protein Stability in Wild-type *E. coli* and the Δ*rfaE* Mutant

Previously, we observed that an increase in stability of LpxC in *E. coli* resulted in a PAE for CHIR-090 (11). LpxC stability is regulated, in part, by LapB and its interaction with PbgA, which modulates LpxC turnover in response to LPS demand (32). Since perturbing RfaE likely alters LPS demand, we hypothesized that the enhanced PAE of LpxC inhibitors in the *rfaE* knockout strain may be driven by reduced LpxC turnover. We therefore investigated the rate of LpxC turnover in the wild-type and Δ*rfaE* strains using pulse SILAC (Stable Isotope Labeling by Amino Acids). We metabolically labelled *E. coli* with isotopically labelled ^13^C_6_ and ^15^N_2_ lysine (Lys8 “heavy lysine”) following growth in media containing normal “light” lysine (Lys0). We then followed the rate of incorporation of “heavy lysine” into LpxC using high-resolution mass spectrometry. After harvesting and lysing the cells, lysates were subjected to LpxC enrichment using an antibody-based pull-down method. The incorporation of the heavy-labelled amino acids in both the wild-type and Δ*rfaE* mutant strain was 100% as determined by the Relative Isotope Abundance (RIA) value at the end of the experiment (33, 34). We measured intervals up to 80 min after transferring to heavy label media and determined that wild-type LpxC has a half-life of 9.06 min, consistent with previously reported values of 4 to 10 min (35–38). In contrast, the rate of LpxC turnover is ∼2-fold slower in the Δ*rfaE* strain (16.88 min) (**Figure 5**, **Tables S2-S4**). As controls, we measured the turnover rates of the housekeeping proteins GAPDH, outer membrane protein A, 30S ribosomal protein 16S, and ribosomal protein S5 in both wild type and the Δ*rfaE* mutant to assess whether protein turnover beyond LpxC was affected by the *rfaE* knockout (**Figure S3, Tables S2-S4**). In general, the turnover rates of the housekeeping proteins showed no significant differences between the two strains, indicating that the modulation of PAE is specific to the genetic knockout of the *rfaE* gene.

**Figure 5:**
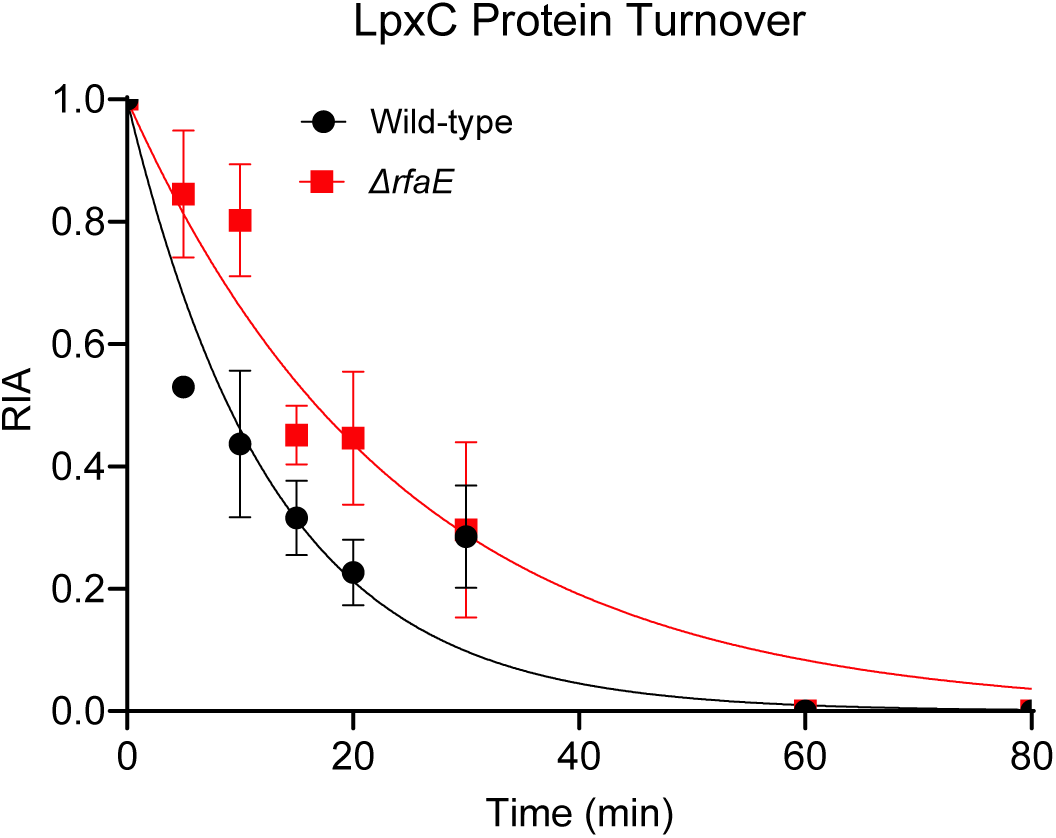
Protein turnover by pSILAC mass spectrometry. Wild-type (black) and Δ*rfaE* (red) protein turnover of LpxC. The relative isotope abundance (RIA) was determined as a function of time after bacteria cultures were transferred to “heavy” lysine-containing media from “light” lysine media. The RIA was calculated by quantifying the isotopic abundance of peptides generated by the trypsin digest using mass spectrometry. Bacteria lysates were enriched using antibody pull-down with beads coated with the LpxC antibody from LSBIO. Experiments were performed independently in triplicate. Data were fit to an exponential decay giving rate constants of 0.078 (95% CI 0.062-0.098) and 0.041 (95% CI 0.035-0.050) min^-1^ for ecLpxC in the wild-type and Δ*rfaE* strains, respectively.

## DISCUSSION

Approaches that increase the PAE of an antibiotic should reduce dosing frequency, widen the therapeutic window and improve patient compliance (2, 4, 5). By decreasing the number of required doses, a prolonged PAE can also limit overall drug exposure, potentially reducing the risk of adverse events, which is a significant barrier to antibiotic approval (39). However, despite the clinical relevance of the PAE, the molecular factors that control this phenomenon are poorly understood. To address this knowledge gap, we developed a method to identify genetic modulators of the PAE using the *E. coli* Keio collection of 3,985 non-essential gene knockouts. We explored this approach with the LpxC inhibitor CHIR-090, which does not produce a PAE in *E. coli* (11).

The PAE is conventionally measured in liquid media; however, this approach is impractical for analyzing genome-scale clone sets. To screen the Keio collection for genetic enhancers of the PAE, we developed a method where bacterial colonies were pinned onto filter paper that enabled each mutant to be simultaneously exposed to the drug of choice. After a wash-out step, the delay in bacterial regrowth was measured as a proxy for the PAE. We initially sought to evaluate this method with CHIR-090 (11) and identified ∼400 gene deletions that enhanced the PAE of this LpxC inhibitor (**Figure 2, Figure S1, Table S1**). We subsequently selected the *rfaE* knockout mutant for further analysis, given the link to LPS biosynthesis and the availability of an RfaE inhibitor, which provided an opportunity to validate the genetic findings with a small molecule. RfaE catalyzes two reactions in the biosynthesis of the LPS inner core precursor ADP-L-glycero-*β*-D-manno-heptose (**Scheme S1**). In the first reaction, D-glycero-D-manno-heptose-7-phosphate is phosphorylated to form D-glycero-*β*-D-manno-heptose-1,7-bisphosphate, after which D-glycero-D-manno-heptose 1-phosphate, formed by the action of the phosphatase GmhB, is coupled with ATP to yield ADP-D-glycero-*β*-D-manno-heptose (14, 23–27). We anticipated differences in PAE values obtained from the primary screen and in liquid media, as bacterial growth on filter paper is monitored by changes in absorbance with a detection limit of 10^5^ CFUs. In contrast, liquid media allows the detection of 10^1^ CFUs (16, 40, 41). In fact, both methods detected increased PAE for CHIR-090 in the *rfaE* knockout. In addition, deleting *rfaE* extended the PAE of LPC-058, suggesting a broader impact on LpxC inhibitors (**Table 1**).

Our long-term goal is to identify drugs that enhance the PAE of antibiotics when used in combination therapy. In this study, co-treatment with an RfaE inhibitor generated a PAE for CHIR-090 and led to a moderate increase in the PAE of LPC-058 (**Table 1**, **Figure 4**). Despite well-documented differences between genetic knockout and chemical inhibition (42), we successfully replicated our genetic findings using a small molecule inhibitor. While previous studies have explored combination therapy to prolong the PAE of existing antibiotics (43, 44), our method uniquely integrates genetic screening with the rational selection of chemical inhibitors that augment the PAE.

We found the enhanced PAE of LpxC inhibitors following perturbation of RfaE was driven by decreased LpxC protein turnover. A similar phenomenon was observed in previous work where chemical and genetic methods were used to reduce the rate of LpxC turnover in *E. coli* (11). Notably, a sub-MIC concentration of the protein synthesis inhibitor azithromycin reduced LpxC turnover by 2-fold, resulting in a PAE of 1.4 – 2.8 h. Here, we demonstrate that deleting *rfaE* leads to a similar decrease in LpxC turnover and a corresponding increase in PAE, reinforcing the link between LpxC stability and PAE duration.

The regulation of LpxC stability is based on the demand for LPS, which is monitored by PbgA and controlled by LapB (**Figure 6**) (32, 45–47). We propose that disrupting RfaE increases the demand for LPS at the outer membrane, thereby promoting its transfer from the inner to the outer membrane. The subsequent depletion of LPS at the inner membrane is sensed by PbgA/LapB, which inhibits FtsH-mediated LpxC degradation, thereby increasing LpxC stability (**Figure 6a**). This regulatory system ensures sufficient LPS synthesis, as higher LpxC stability enhances LPS production. However, increased stability also makes the enzyme more vulnerable to inhibition. Specifically, the increase in stability likely prolongs drug-target residence time between LpxC inhibitors and LpxC, enhancing the PAE. Conversely, under normal conditions where RfaE is unperturbed, more rapid LpxC turnover leads to relatively drug-free target and a reduced or absent PAE.

**Figure 6:**
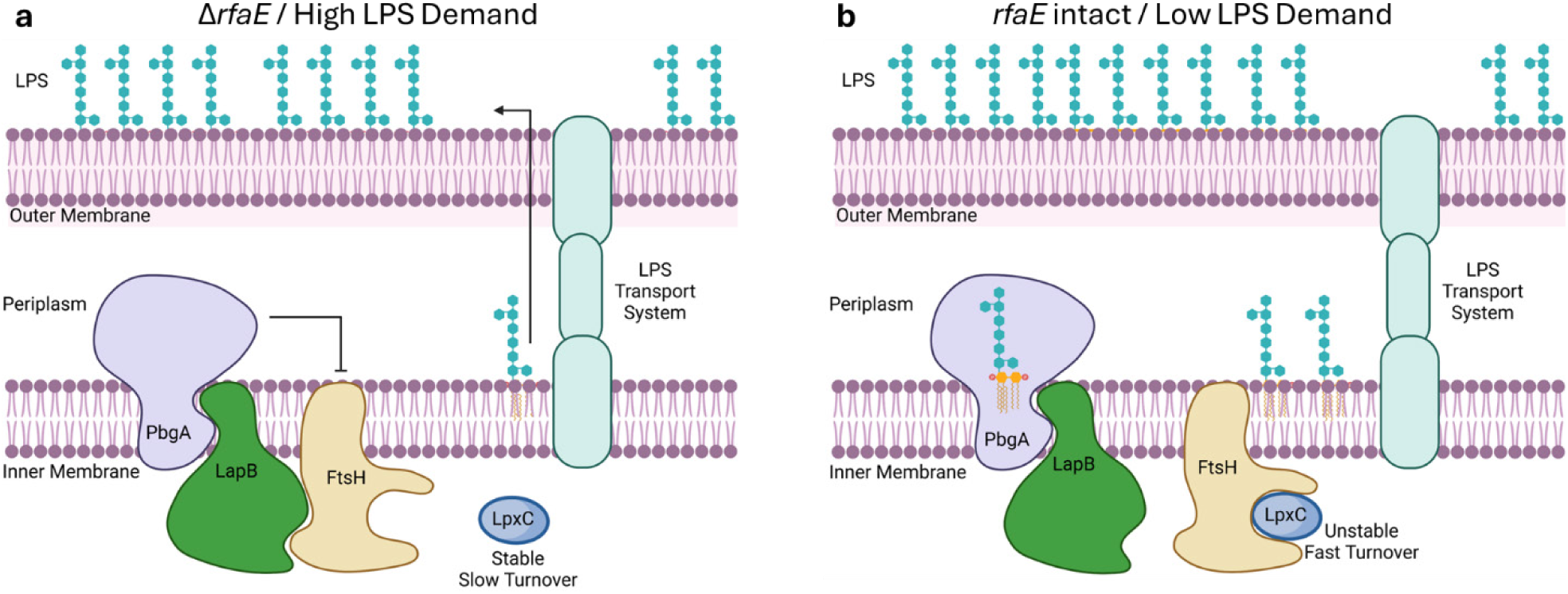
PbgA detects periplasmic LPS levels to regulate LpxC stability through FtsH-mediated degradation. (a) When *rfaE* is knocked out, the demand for LPS in the outer membrane increases. This reduces the level of LPS in the periplasm inner leaflet, which causes PbgA-LapB to antagonize FtsH leading to an increase in LpxC stability, an increase in production of the LPS precursor Lipid A and hence increased LPS synthesis. (b) When sufficient LPS is present in the outer membrane, excess LPS formed in the periplasm inner leaflet binds to PbgA preventing the PbgA-LapB complex from antagonizing FtsH which promotes the degradation of LpxC.

## Conclusion

Strategies that extend the PAEs of antibiotics can improve drug safety and patient compliance by reducing dosing frequency and widening the therapeutic window. In the present work, we developed a chemical-genetic screen to identify modulators of the PAE and applied it to the LpxC inhibitor CHIR-090, which does not generate a PAE in *E. coli*. We identified ∼400 deletions that increased the PAE of CHIR-090 and subsequently focused on the Δ*rfaE*, a gene involved in the biosynthesis of the LPS core oligosaccharide. We propose that deletion of *rfaE* or chemical inhibition of RfaE increases the demand for LPS, leading to a 2-fold decrease in the rate of LpxC turnover. The resulting increase in LpxC stability extends the PAE for CHIR-090 and the LpxC inhibitor LPC-058 by 2-4-fold, reinforcing the direct link between protein synthesis and time-dependent drug activity. More broadly, this study provides a foundation for discovering unexplored targets that enhance the PAE through system-wide genetic perturbation.

## MATERIALS AND METHODS

### Reagents

CHIR-090 was purchased from Fisher Scientific, and L-Lysine:2HCl (13C6,99%, 15N4,99 from Cambridge Isotope Laboratories (Fisher Scientific). The LpxC antibody was purchased from LSBIO. The synthesis of **1** is described in the supplementary information using synthetic routes adapted from Desroy et al. (15). The synthesis of LPC-058 is described in the supplementary information using the synthetic routes adapted from the original publications of the compound (31, 48).

### Bacterial Strains

The Keio collection was originally developed by Dr. Mori at the Keio University in Japan (12). Both the Keio collection and the K-12 derivative wild-type strain BW25113 were available at McMaster University (16). The *E. coli ΔacrAB-ΔtolC* strain was a kind gift from Professor Zgurskaya at the University of Oklahoma.

### Minimum Inhibitory Concentration (MIC) Measurements on Solid Media

The Keio collection was grown from frozen stocks on Luria-Bertani (LB) agar media containing 50 µg/mL of kanamycin at 96-well density. Colonies were grown overnight at 37°C, then scaled up to 384-density and 1536-density using a Singer Rotor (Singer Instruments, Somerset, UK). The 384-density and 1536-density plates were duplicated on LB media to make master plates, which were maintained at 4°C for 2-3 weeks.

MIC values for CHIR-090 were determined as previously described (16). Briefly, an initial bed of 25 mL of 2% agarose was poured into an empty Singer PlusPlate and allowed to dry. A size 16 test tube was used to cut holes in the agarose, and each hole was filled with LB containing different concentrations of the drug until it was perfectly level with the agarose (∼440 mL). The LA media plugs were allowed to dry, and the agarose template was removed, leaving pads of medium containing a concentration gradient of the drug. *Escherichia coli* K-12 BW25113 was pinned at 384 density from a master plate onto the agar plugs and was grown for 18 h at 37°C.

### Minimum Inhibitory Concentration (MIC) Measurements in Liquid Culture

Antibacterial susceptibility tests for aerobically grown bacteria were performed using the micro broth dilution assay in transparent round-bottom 96-well plates where bacterial growth was assessed by visual inspection in accordance to the Clinical and Laboratory Standard Institute (49). Bacteria were grown to mid log phase (OD_600_ of 0.4-0.6) in cation-adjusted Mueller-Hinton (CaMH) media at 37°C in an orbital shaker. A final concentration of 10^6^ CFU/mL per well was added to media containing 2-fold dilutions of compounds, which generated final concentrations of compounds ranging from 0.009 to 100 μM. The MIC was the concentration of compound where no visible growth could be seen after 18 h incubation at 37°C.

A checkerboard assay was performed to determine the MIC values of erythromycin, CHIR-090, and LPC-058 in the presence of **1**. Concentrations of erythromycin, CHIR-090, and LPC-058 ranged from 0-100 μM on one axis, while **1** varied from 0-100 μM on the second axis. Wells with no growth were determined by visual inspection. All MICs were determined in triplicate and validated in triplicate through standard MIC protocols.

### Chemical Genetic Screening

Chemical-genetic screens were performed by pinning the Keio collection in 1536-density onto LB agar supplemented with CHIR-090. The media was supplemented with 0.015 µg/mL (1/8x MIC), 0.03µg/mL (1/4x MIC), 0.06 µg/mL (1/2x MIC), or 0.25 µg/mL (2x MIC) of CHIR-090. Cells were grown at 37°C for 18 h on LB containing CHIR-090. After incubation, plates were scanned in transmissive mode on an Epson Perfection V750-M scanner and quantitatively analyzed using ImageJ as previously described (50). Edge effects were normalized using a double-pass method across columns and rows based on the median value. Strains grown at 37°C in the presence of CHIR-090 with a lag time of 30 h compared to the no-drug were defined as sensitive.

### Post-Antibiotic Effect (PAE) Lag Time on Solid Media

The Keio collection was pinned onto 8 x 12 cm nitrocellulose membranes at 384 density, which had been placed onto solid LB agar supplemented with 50 µg/mL of kanamycin in PlusPlates (Singer Instruments). Plates were incubated at 37°C for 16 h. The membranes were then transferred with sterile forceps onto LB agar or LB agar supplemented with 2 µg/mL (16x MIC) of CHIR-090. After incubation for an additional 6 h at 37°C, 384-long pins were used to manually pin the colonies into a 384-well plate containing 50 µL of PBS per well and then transferred to LB agar plates containing 50 µg/mL of kanamycin using the Singer Rotor. Plates were then incubated at 37°C for at least 10 h and scanned every 20 min using the Epson Perfection V750 scanners after which the integrated densities were extracted for each colony.(16, 17)

The lag time of each colony was calculated using an in-house script written in R. The change in PAE for each mutant was determined by normalizing the lag time of each mutant after exposure to the drug to the interquartile mean of the plate, followed by normalization to the no-drug control condition. The change in PAE was calculated using equation **1**, where the lag time of the mutant is LT_m_, and the interquartile mean of the respective plate is LT_IQM._ A

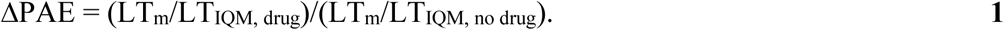

Experiments were conducted in quadruplicate, and strains with a ΔPAE of at least one standard deviation less than the mean were considered enhancers of the PAE.

### Post-antibiotic Effect (PAE) in Liquid Culture

Bacteria were grown to mid-log phase (OD_600_ of 0.4-0.6) in CaMH media at 37°C, followed by exposure to varying concentrations of vehicle (DMSO or nuclease-free water), CHIR-090, or LPC-058. After shaking for 30 to 60 min on an orbital shaker at 37°C, cultures were diluted 1000-fold into fresh CaMH media, and the regrowth of bacteria was monitored by taking 100 μL aliquots at 1 h time intervals. Serial dilutions were plated on Muller-Hinton agar plates, which were incubated overnight (16-18 h) at 37°C after which CFUs were determined by counting colonies. The PAE was calculated as the time needed for the bacteria to regrow 1 log_10_ CFU subtracted from the time needed for the control (vehicle, no drug, 0x) to increase by 1 log_10_ CFU.(1) Each time point was performed at a minimum in duplicate and at a maximum in quintuplet, and the entire PAE experiment was performed in duplicates (n=2) or triplicates (n=3).

### Time-Kill Assays

Bacteria were grown to mid-log phase (OD_600_ of 0.4-0.6) in CaMH media at 37°C, followed by exposure to a range of 1x up to 16x MIC concentration of compound or vehicle (DMSO, nuclease-free water). During the exposure period, 100 μL aliquots were taken at 15 min intervals up to 60 or 90 min and serially diluted 10-fold 5 times generating a dilution range of 0-100,000-fold. Ten μL of each serial dilution in a 96-well plate was then plated at the top of Muller-Hinton agar plates, which was then tilted to allow the culture to travel down the plate. Before the culture reached the bottom of the plate, the plate was tilted in the opposite direction to allow the culture to travel back up toward the original deposit of the culture, which gave a semi-even distribution of liquid. The CFUs were then determined by counting colonies after incubating at 37°C overnight (16-18 h).

### pSILAC

The *E. coli* wild-type (BW25113) and the Δ*rfaE* strains were first grown overnight in 5 mL CaMH media. The media was then replaced with 100 mL of M9 Minimal Media (Sigma-Aldrich) consisting of 11.28 g/L M9 salts, 5g/L glucose, 1 mM MgS0_4_, 0.1 mM CaCl_2_, 33 μM thiamine, 1 mM trace metal solution, and 200 mg/L of the 20 essential amino acids, containing the “light” isotopes of each amino acid. The cells were grown at 37°C in an orbital shaker at 250 rpm until an OD_600_ of 0.4-0.6 was reached. A 5 mL aliquot was removed at time 0, and the cells were then harvested by centrifugation at 3900 rpm for 20 min. The remaining culture was also pelleted, followed by washing 3 times with 1x Hank’s Blank Salt Solution (HBSS, Sigma Aldrich), transferred to M9 media containing 200 mg/L “heavy” L-Lysine:2HCL (13C6,99%, 15N4,99) together with the remaining 19 essential amino acids, then incubated at 37°C in the shaking incubator at 250 rpm. Aliquots (5 mL) of the culture were taken at 0, 5, 10, 15, 20, 30, 60, and 80 min, and the cells were pelleted by centrifugation at 3900 rpm for 20 min. Pellets were resuspended in 1x HBSS and centrifuged at 3900 rpm for 20 min.

The pellets were lysed in 500 μL lysis buffer containing 1% Triton X-100, PBS pH 7.4, and protease inhibitor cocktail buffer by sonication at level 3 for 10 s followed by recovery on ice for 5 min. The sonication was repeated two more times. Cell debris was removed by centrifugation at 13,000 rpm for 45 min at 4°C, and 400 μL of the lysate was then transferred to a sterile microcentrifuge tube. Total protein concentration was calculated by the BCA assay (Pierce BCA Kit, Thermo Scientific), and lysis buffer was added to give 300 μg protein in a volume of 500 μL. LpxC was then immunoprecipitated by adding 5 μg of LpxC antibody (LS Bio), incubating for 2 h at RT, followed by adding 50 μL of Pierce Magnetic G beads (Fisher Scientific) that had been pre-washed in lysis buffer. After 1 h, the beads were isolated and washed with 500 μL of lysis buffer three times. Subsequently, the beads were exposed to 12 mM iodoacetamide for 45 min at RT to alkylate the cysteines and then digested with trypsin at 37°C (1:1000) overnight. Peptides were then desalted on C18 S-Trap cartridges, lyophilized and resuspended in 0.1% formic acid before LC-MS/MS analysis. Experiments were performed independently in triplicate.

### Peptide Identification and Quantification by LC/MS

Peptides were analyzed by nano LC-MS/MS. Parent peptide mass, collision-induced fragment mass information, and isotopically encoded peptide abundance values were obtained by liquid chromatography-electrospray ionization tandem mass spectrometry (LC-MS/MS) using an orbital trap instrument (Thermo Q-Exactive HF) followed by protein database searching against a target database of *E. coli* K12. HPLC C18 columns (75 μm ID × ∼15 cm) and were self-packed with 3 μm Reprosil C 18 resin. Peptides were separated on the resolving column with a flow rate of 300 nL/min using a gradient elution step 0−40% with acetonitrile (MeCN) and 0.1% formic acid (0.23%/min) over 40 min followed by a 10 min wash with 90% MeCN and a 10 min wash step with isocratic 90% MeCN. Electrospray ionization was achieved using a spray voltage of ∼2.3 kV. Data-dependent MS and MS-MS acquisitions were made using a survey scan (m/z 400−1600) with a maximum fill of 50 ms followed typically by 20 consecutive product ion scans (m/z 100−1800). Parent ion with charge states of 2+, 3+, and 4+ were selected with a 15 s exclusion period. MS data were collected using Xcaliber (Thermo).

Proteins were identified from the survey and product ion spectra data using Proteome Discoverer 2.4 (Thermo). Two missed tryptic cleavages were allowed, and posttranslational modifications were considered including cysteine carbamidomethylation, NQ-deamidation, and ST-dehydration. Quantitation was based on heavy/light Lys incorporation. Database searches used *E. coli* UniProt FASTA database (including common contaminants). False discovery rates of protein identification were binned at <1% and <5%.

### Protein Turnover Calculation

The protein turnover calculation was performed as described.(51, 52) Briefly, the relative isotopic abundance at time t (RIA_t_) was calculated by dividing the abundance of light isotope (AL) by the sum of the abundance of both light (AL) and heavy isotopes (AH) (equation **2**).

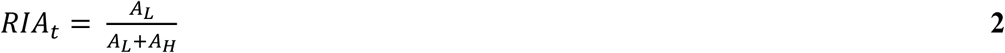

The data obtained from SILAC were normalized and fit to a one-phase exponential decay equation in GraphPad Prism 9 (equation **3**),

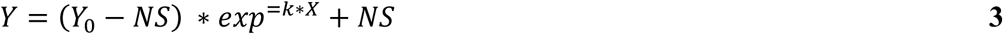

where Y is RIA_t_ (RIA at specific time point t), Y_0_ is RIA at t = 0 (RIA_0_), NS is RIA at t ∞ (RIA_∞_, t_max_ 80 min; Plateau), k is k_loss_, and X is time (t). The above equation can be related to equation **4**,

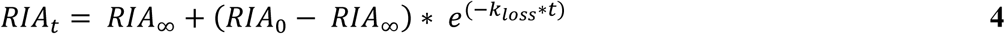

where RIA_∞_ is the relative isotope abundance at infinite time after starting the experiment, and RIA_0_ is the relative isotope abundance at the start of the chase experiment. The half-life was calculated by dividing 0.693 by k_loss_.

## AUTHOR INFORMATION

## Notes

The authors declare no competing financial interest.

## AUTHOR CONTRIBUTIONS

Conceptualization, A.L.G., M.M.T., K.R., E.D.B. and P.J.T.; methodology A.L.G., M.M.T., K.R., S.G.W., J.D.H., E.D.B. and P.J.T.; investigation A.L.G., M.M.T., K.R., F.A., M.A., M.I.A., S.A.C., P.D.., A.C.P.K., A.S., J.T.C. and J.D.H.; formal analysis, A.L.G., M.M.T., K.R., J.D.H.; writing – original draft, A.L.G. and P.J.T.; writing – review and editing, A.L.G., M.M.T., E.D.B. and P.J.T.; funding acquisition, E.D.B. and P.J.T.; resources, E.D.B. and P.J.T.; supervisor, P.J.T.

## ACKNOWLEDGMENTS

This study was supported by the National Institutes of Health grant R35GM149297 to P.J.T, in addition to a Tier 1 Canada Research Chair award, a Foundation Grant from the Canadian Institutes of Health Research (CHIR; FRN 143215) and a grant from the Ontario Research Fund (RE09-047) to E.D.B. A.L.G. was supported by the National Institutes of Health Chemistry-Biology Interface Training Grant (T32GM092714). S.A.C. was supported by the National Institute of Health Scholars in Biomedical Sciences Training Grant (T32GM127253). M.M.T. was supported by a CIHR Canada Graduate Scholarship (CGS-D). J.T.C. was supported by the National Institutes of Health IMSD-MERGE (T32GM135746) Program at Stony Brook University. A.S. and J.T.C. were supported by the NY-CAPs IRACDA (K12-GM102778) Program at Stony Brook University.

