## Supplementary Information for "Identifying Modulators of the Post-Antibiotic Effect"

**Table of Contents**

Figure S1: Lag time of selected Keio strains on solid media following exposure to CHIR-090. S3

Figure S2. Time-kill experiments of CHIR-090 and LPC-058 in the ΔrfaE mutant. .................. S4

Figure S3: Protein Turnover by pSILAC mass spectrometry. ................................................... S5

Table S1. Microbiological data for selected Keio strains on solid media with CHIR-090 .......... S6

Table S2: Proteins Analyzed by SILAC from the *E. coli* strain BW25113 ................................ S7

Table S3: Proteins Analyzed by SILAC from the *E. coli* strain Δ*rfaE* ....................................... S8

Table S4: Protein Turnover Rates from pSILAC Mass Spectrometry ........................................ S9

Scheme S1: Biosynthesis of ADP-L-*glycero*-B-D-*manno*-heptose ........................................... S10

General synthesis methods ........................................................................................................ S11

Synthesis of RfaE inhibitor **1** ..................................................................................................... S12

Scheme S2. Synthetic scheme for the RfaE inhibitor **1** ............................................................ S12

Synthesis of LPC-058 .................................................................................................................. S17

Scheme S3: Synthesis Scheme for the LpxC inhibitor, LPC-058. ............................................ S17

References S21

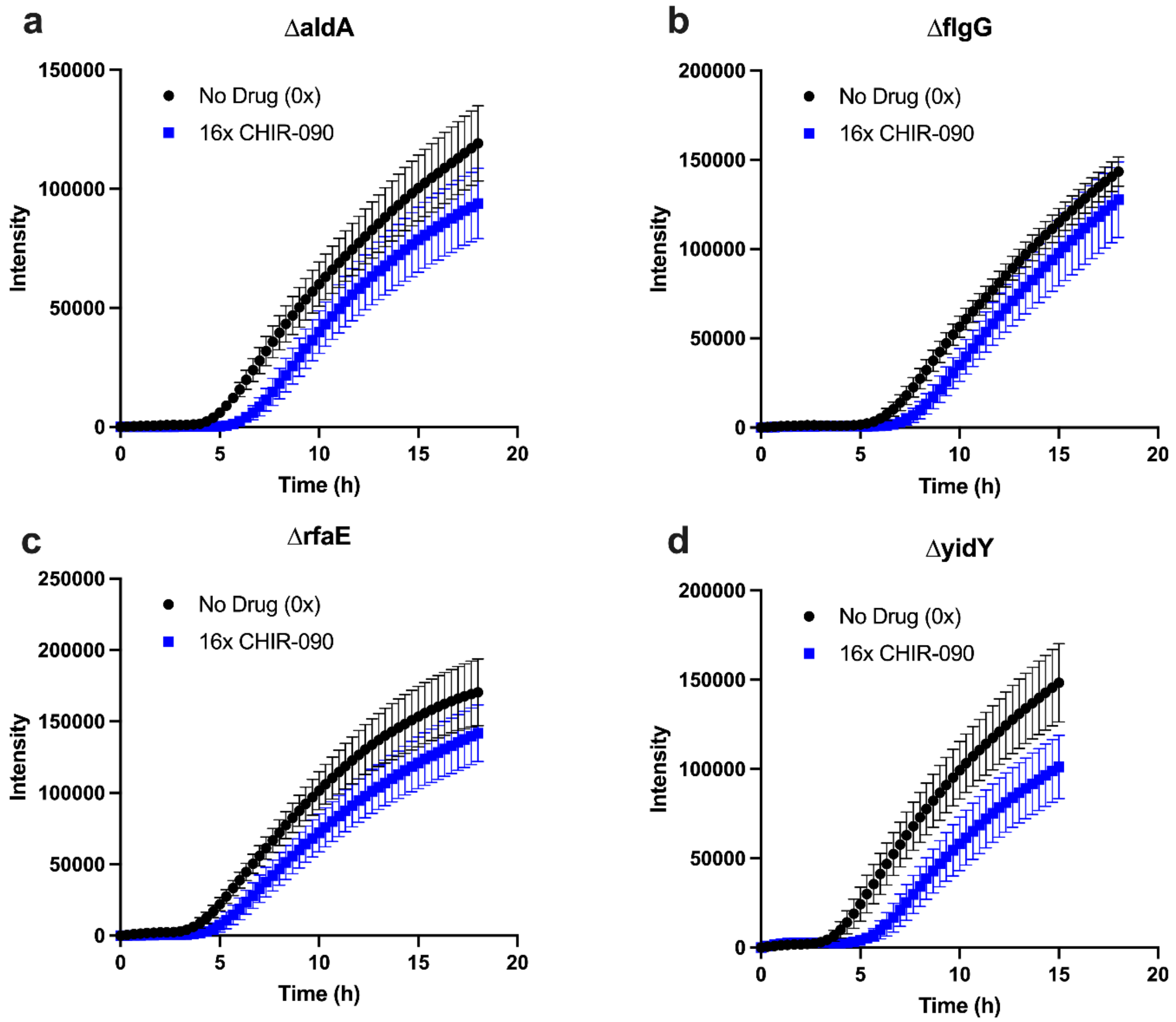


**Figure S1: Lag time of selected Keio strains on solid media following exposure to CHIR-090.** These confirmatory studies employed growth on solid media as described for the primary screen. Shown are the wild-type strain K-12, BW25113, and Δ*aldA*, Δ*flgG*, Δ*rfaE*, and Δ*yidY*. Regrowth of the bacterial colonies is shown in black for no drug treatment and blue for 16x MIC CHIR-090.

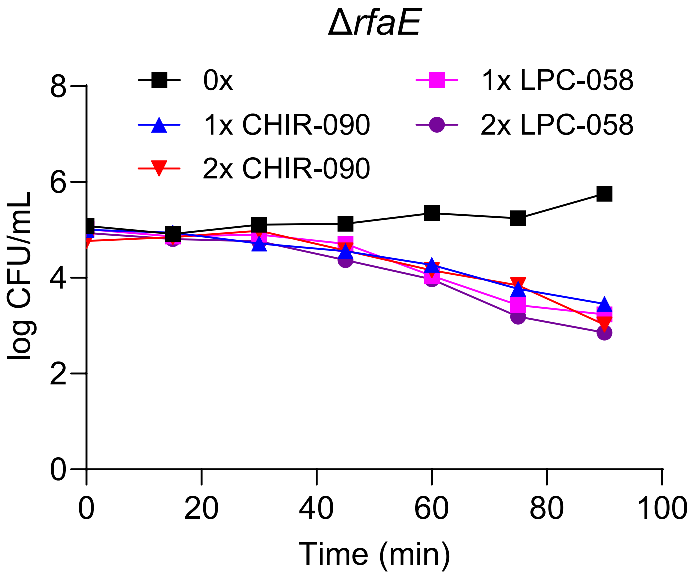


**Figure S2. Time-kill experiments of CHIR-090 and LPC-058 in the ΔrfaE mutant.** Cultures of bacteria ($10^6$ CFU/mL) were grown to mid-log phase (OD 600 of 0.6-0.7) in CaMH media at 37°C and then exposed to 1x and 2x MIC of CHIR-090 or LPC-058 or vehicle 0x (DMSO). Subsequently, 0.1 mL aliquots were withdrawn at 15, 30, or 60-minute time intervals and plated in 10 μL serial dilutions on Muller-Hinton agar plates. The CFUs were determined by counting colonies after overnight incubation at 37°C. Experiments were done in duplicate.

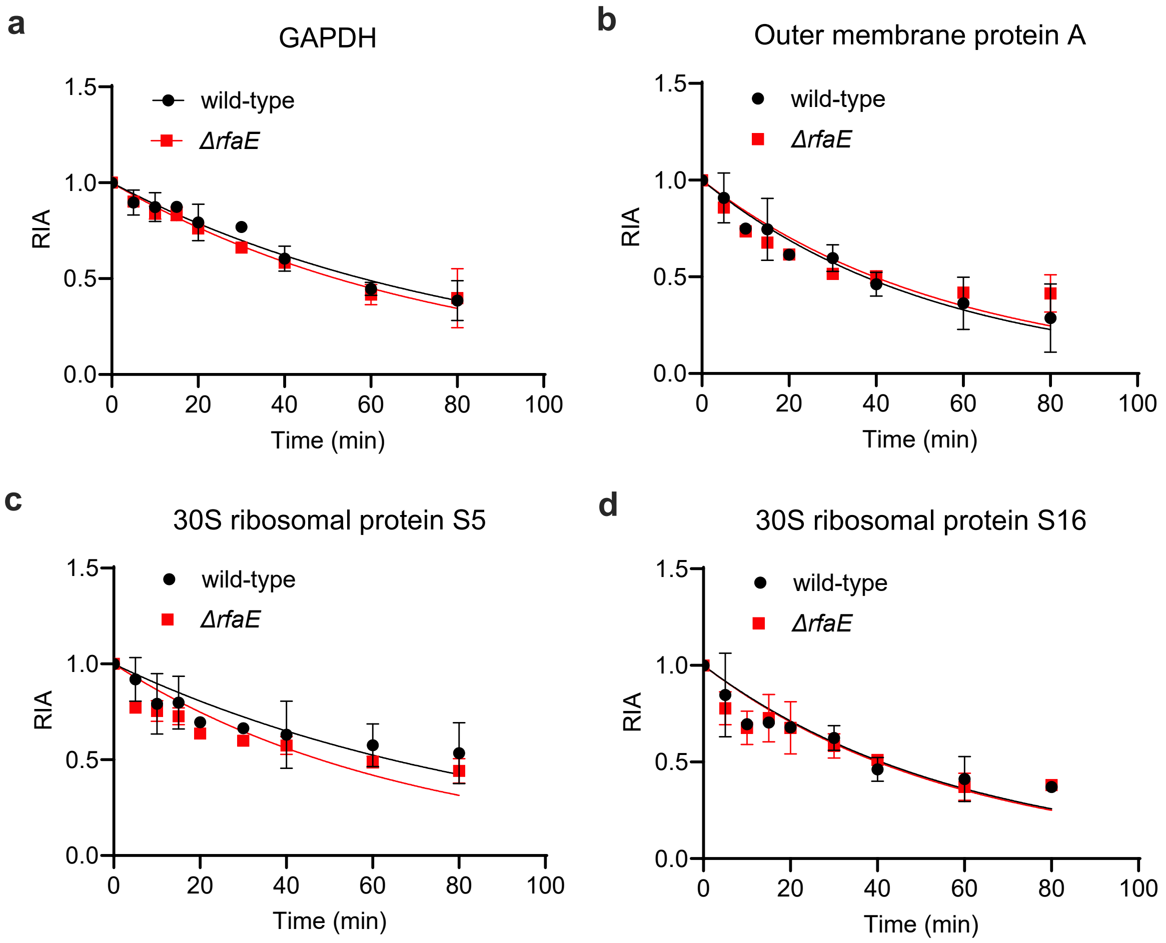


**Figure S3: Protein Turnover by pSILAC mass spectrometry.** (a) GAPDH, (b) outer membrane protein A, (c) 30S ribosomal protein S1 and (d) the 30S ribosomal S16 in *E. coli.* The relative isotope abundance (RIA) was determined as a function of time after bacteria cultures were transferred to "heavy" lysine-containing media from "light" lysine media. The RIA was calculated by quantifying the isotopic abundance of peptides generated by the trypsin digest using mass spectrometry. Bacteria lysates were enriched using antibody pull-down with beads coated with the LpxC antibody from LSBIO.

| Table S1. Microbiological data for selected Keio strains on solid media with CHIR-090 | | |
|---|---|---|
| | CHIR-090 | |
| Strain | MIC (μM)[b] | Δlag time (h)[c] |
| WT[a] | 0.29 | 0 |
| Δ*aldA* | 0.29 | 1.67 |
| Δ*flgG* | 0.29 | 1.33 |
| Δ*rfaE* | 0.07 | 1.1 |
| Δ*yidY* | 0.29 | 1.91 |

[a] WT, wild-type, *E. coli* K-12, BW25113.

[b]MIC values were determined by plating the Keio collection on 1536-colony plates treated with sub- and supra-MIC concentrations of CHIR-090 based on the MIC for the wild-type strain to determine sensitivity to the drug. Experiments were performed in triplicate.

[c]Δlag time is the difference in the time taken for the treated bacteria to start growing compared to the untreated strain. This is assumed to represent the PAE in solid media. Lag time plates were tested in quadruplicate.

**Table S2: Proteins Analyzed by SILAC from the *E. coli* strain BW25113**

| Accession Number | Gene | Unique Peptides | | PSMs | | MW (kDa) | SUM PEP Score | | RIA Values (L/L+H) | | | | | | | |
|---|---|---|---|---|---|---|---|---|---|---|---|---|---|---|---|---|
| | | E1 | E2 | E1 | E2 | | E1 | E2 | 0 | 5 | 10 | 15 | 20 | 30 | 60 | 80 |
| P0A725 | *lpxC* | 15 | 17 | 191 | 88.79 | 33.9 | 54.65 | 256 | 1.0 ±0.0 | 0.53 ±0.009 | 0.44 ±0.12 | 0.32 ±0.06 | 0.23 ±0.05 | 0.29 ±0.08 | 0.0 ±0.0 | 0.0 ±0.0 |
| P0A9B2 | *gapA* | 23 | 26 | 942 | 6259 | 35.5 | 206.77 | 447.33 | 1.0 ±0.001 | 0.90 ±0.07 | 0.87 ±0.08 | 0.87 ±0.03 | 0.79 ±0.09 | 0.77 ±0.0001 | 0.45 ±0.03 | 0.39 ±0.10 |
| P0A910 | *ompA* | 13 | 2 | 191 | 3114 | 37.2 | 63.93 | 295.75 | 1.0 ±0.008 | 0.91 ±0.13 | 0.75 ±0.03 | 0.75 ±0.16 | 0.62 ±0.002 | 0.60 ±0.07 | 0.36 ±0.14 | 0.29 ±0.18 |
| P0A7T3 | *rpsP* | 7 | 5 | 126 | 848 | 9.2 | 43.73 | 97.14 | 1.0 ±0.004 | 0.85 ±0.22 | 0.70 ±0.02 | 0.71 ±0.003 | 0.68 ±0.01 | 0.62 ±0.06 | 0.41 ±0.12 | 0.37 ±0.03 |
| P0A7W3 | *rpsA* | 36 | 40 | 916 | 6155 | 61.1 | 261.77 | 656.84 | 1.0 ±0.001 | 0.92 ±0.11 | 0.79 ±0.16 | 0.80 ±0.14 | 0.70 ±0.02 | 0.66 ±0.03 | 0.58 ±0.11 | 0.53 ±0.16 |

Unique peptides: Defined as a peptide that exists in only one protein in the proteome of interest (1).

PSMs: Scoring function that assigns a numerical value to the peptide-spectrum (P-S) pair expressing the likelihood that the fragmentation of a peptide with sequence P is recorded in experimental mass spectrum S. PSM score informs on the quality of the spectrum (2).

Sum PEP Score: Score that signifies the probability of the observed PSM being incorrect (3). Scores are reported as -10*log10(P), where P is the absolute probability. A probability of 10-20 thus becomes a score of 200.

RIA Values (H/H+L): Relative Isotopic Abundance. Abundance of heavy labelled peptide (H), divided by the abundance of all (heavy+ light) peptide.

Proteins, OS=*Escherichia coli* (strain K12)

*lpxC*: UDP-3-*O*-acyl-*N*-acetylglucosamine deacetylase; *gapA*: Glyceraldehyde-3-phosphate dehydrogenase A; *ompA*: Outer membrane protein A; *rpsP*: 30S ribosomal protein S16; *rpsA*: 30S ribosomal protein S5.

**Table S3: Proteins Analyzed by SILAC from the *E. coli* strain Δ*rfaE***

| Accession Number | Gene | Unique Peptides | | PSMs | | MW (kDa) | SUM PEP Score | | RIA Values (L/L+H) | | | | | | | |
|---|---|---|---|---|---|---|---|---|---|---|---|---|---|---|---|---|
| | | E1 | E2 | E1 | E2 | | E1 | E2 | 0 | 5 | 10 | 15 | 20 | 30 | 60 | 80 |
| P0A725 | *lpxC* | 9 | 17 | 57 | 246 | 33.9 | 42.6 | 88.79 | 1.0<br>±0.0 | 0.85<br>±0.10 | 0.81<br>±0.09 | 0.45<br>±0.05 | 0.45<br>±0.11 | 0.30<br>±0.14 | 0.0<br>±0.0 | 0.0<br>±0.0 |
| P0A9B2 | *gapA* | 28 | 30 | 13144 | 1810 | 35.5 | 555.9 | 302.41 | 1.0<br>±0.0 | 0.90<br>±0.01 | 0.85<br>±0.03 | 0.84<br>±0.04 | 0.76<br>±0.003 | 0.66<br>±0.02 | 0.42<br>±0.05 | 0.40<br>±0.15 |
| P0A910 | *ompA* | 4 | 13 | 7353 | 178 | 37.2 | 473.78 | 72.92 | 1.0<br>±0.007 | 0.86<br>±0.01 | 0.74<br>±0.01 | 0.68<br>±0.02 | 0.62<br>±0.02 | 0.52<br>±0.03 | 0.42<br>±0.005 | 0.41<br>±0.01 |
| P0A7T3 | *rpsP* | 7 | 7 | 1662 | 238 | 9.2 | 133.66 | 71.08 | 1.0<br>±0.007 | 0.78<br>±0.09 | 0.68<br>±0.09 | 0.73<br>±0.12 | 0.68<br>±0.14 | 0.58<br>±0.06 | 0.37<br>±0.07 | 0.38<br>±0.01 |
| P0A7W3 | *rpsA* | 11 | 10 | 3288 | 612 | 61.1 | 404.98 | 153.23 | 1.0<br>±0.0 | 0.77<br>±0.006 | 0.75<br>±0.05 | 0.73<br>±0.05 | 0.64<br>±0.03 | 0.60<br>±0.02 | 0.49<br>±0.03 | 0.44<br>±0.06 |

Unique peptides: Defined as a peptide that exists in only one protein in the proteome of interest (1).

PSMs: Scoring function that assigns a numerical value to the peptide-spectrum (P-S) pair expressing the likelihood that the fragmentation of a peptide with sequence P is recorded in experimental mass spectrum S. PSM score informs on the quality of the spectrum (2).

Sum PEP Score: Score that signifies the probability of the observed PSM being incorrect (3). Scores are reported as -10*log10(P), where P is the absolute probability. A probability of 10-20 thus becomes a score of 200.

RIA Values (H/H+L): Relative Isotopic Abundance. Abundance of heavy labelled peptide (H), divided by the abundance of all (heavy+ light) peptide.

Proteins, OS=*Escherichia coli* (strain K12)

lpxC: UDP-3-*O*-acyl-*N*-acetylglucosamine deacetylase; gapA: Glyceraldehyde-3-phosphate dehydrogenase A; ompA: Outer membrane protein A; rpsP: 30S ribosomal protein S16; rpsA: 30S ribosomal protein S5.

**Table S4: Protein Turnover Rates from pSILAC Mass Spectrometry**

| | **Wild-type** | | **Δ*rfaE*** | |
|---|---|---|---|---|
| **Protein** | k ($min^{-1}$) | Half-life (min) | k ($min^{-1}$) | Half-life (min) |
| LpxC | 0.077 (0.061 to 0.097) | 9.06 (7.12 to 11.38) | 0.041 (0.034 to 0.049) | 16.88 (14.03 to 20.32) |
| Glyceraldehyde-3-phosphate dehydrogenase | 0.012 (0.010 to 0.013) | 59.17 (51.63 to 68.31) | 0.013 (0.012 to 0.015) | 52.53 (46.56 to 59.55) |
| Outer membrane protein A | 0.018 (0.015 to 0.022) | 38.14 (30.94 to 47.30) | 0.017 (0.014 to 0.022) | 39.80 (31.92 to 49.94) |
| 30S Ribosomal protein S16 | 0.016 (0.013 to 0.021) | 42.47 (33.54 to 54.32) | 0.017 (0.013 to 0.022) | 39.75 (30.74 to 51.89) |
| 30S Ribosomal protein S5 | 0.011 (0.008 to 0.014) | 65.51 (49.52 to 89.69) | 0.015 (0.011 to 0.019) | 47.41 (36.80 to 61.97) |

Turnover SILAC experiments were performed in M9 media with wild-type (BW25113) and the Δ*rfaE E. coli* strains. Experiments were performed in duplicate and data were fit to a single exponential decay to obtain the rate of protein turnover. Numbers in parentheses are the 95% confidence intervals.

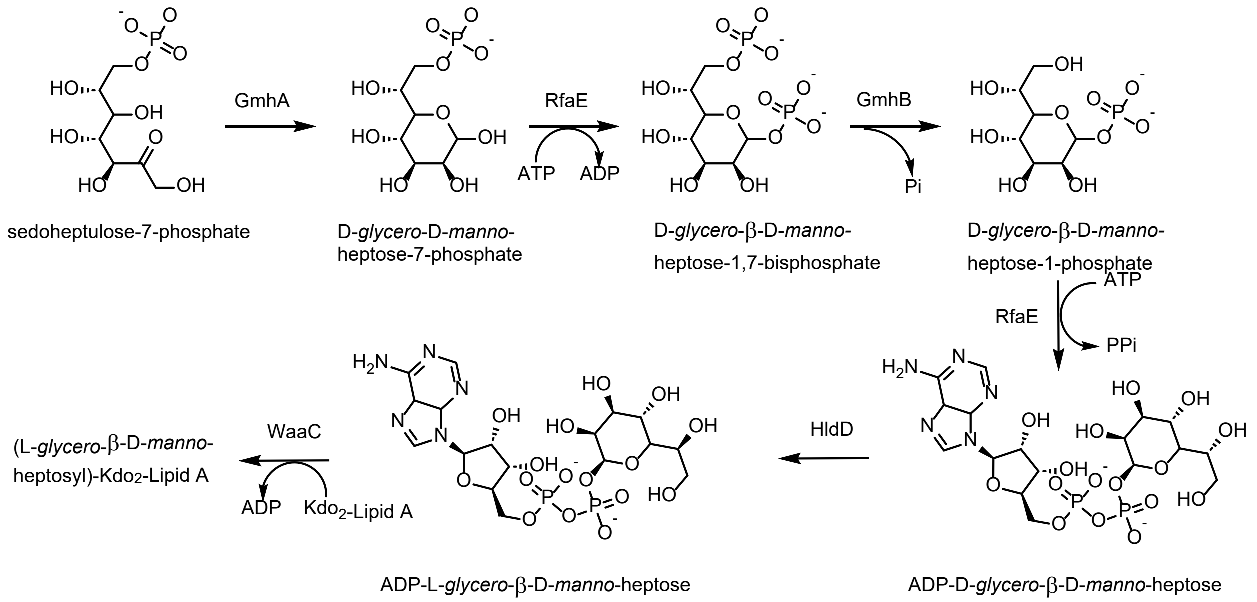


**Scheme S1: Biosynthesis of ADP-L-*glycero-B*-D-*manno*-heptose.**

**General synthesis methods**

All commercial chemicals and reagents were used as received unless otherwise noted. Crude products were purified via column chromatography using a Teledyne ISCO Combiflash NextGen 100 system. $^1$H nuclear magnetic resonance spectroscopy (NMR) and $^{13}$C NMR spectra were recorded on a Bruker Avance 500 MHz spectrometer. Chemical shifts are expressed in δ ppm referenced to the residual solvent peak ($CDCl_3$, δ = 7.26 ppm; $d_6$-DMSO, δ = 2.50 ppm). Abbreviations used in describing peak signals are as follows: br, broad signal; v, very; s, singlet; d, doublet; dd, doublet of doublets; t, triplet; q, quartet; m, multiplet. Low-resolution mass spectra were recorded on an Agilent 6100 series single quadrupole mass spectrometer equipped with an Agilent 1290 Infinity II high performance liquid chromatography system. High-resolution mass spectra were recorded on a Bruker Impact II quadrupole time-of-flight mass spectrometer (QTOF MS) equipped with an Agilent 1290 Infinity II high performance liquid chromatography system.

### Synthesis of RfaE inhibitor 1.

The synthesis of **1** was adapted from Desroy *et al* (4).

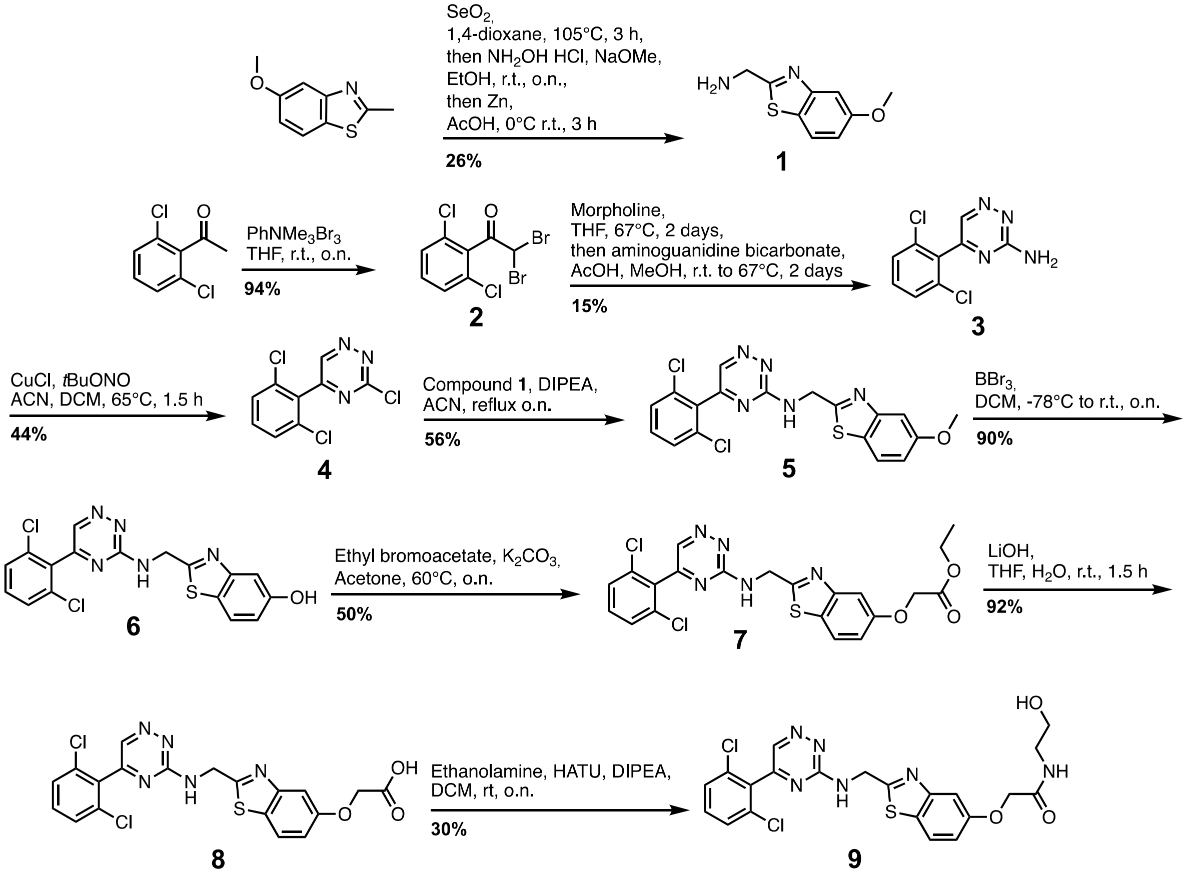


**Scheme S2. Synthetic scheme for the RfaE inhibitor 1.**

### Experimental details for Scheme S2.

#### (5-Methoxy-benzothiazol-2-yl)-methylamine (1)

Under argon, selenium dioxide (0.929 g, 8.37 mmol) was added to a solution of 5-methoxy-2- methyl-1,3-benzothiazole (1.000 g, 5.58 mmol) in 1,4-dioxane (15 mL). The reaction mixture was stirred at 105 °C for 3 hours, then filtered over a pad of celite and rinsed with hot 1,4-dioxane. The filtrate was concentrated to give crude 5-methoxy-1,3-benzothiazole-2-carbaldehyde. The latter compound was dissolved in ethanol (12 mL), hydroxylamine hydrochloride (0.776 g, 11.16 mmol) and sodium methylate (0.714 g, 13.22 mmol) were successively added and the reaction mixture was stirred at room temperature overnight. The solution was filtered and concentrated to give crude 5-methoxy- 1,3-benzothiazole-2-carbaldehyde oxime. The latter compound was dissolved in acetic acid (30 mL), and zinc dust (2.189 g, 33.48 mmol) was added portionwise. The solution was stirred at room temperature for 3 hours. The mixture was then filtered over a pad of celite and rinsed with methanol. Toluene was added and the filtrate was concentrated. Water was added, then the solution was acidified to pH = 2 with hydrochloric acid 1N and extracted with ethyl acetate. These organic layers were discarded. The aqueous layer was basified with aqueous sodium hydroxide to pH = 12, and then extracted with ethyl acetate. Combined organic extracts were dried over magnesium sulfate, filtered and evaporated to give (5-methoxy-1,3-benzothiazol-2-yl)methylamine **1** as an orange solid (0.783 g, 72%) to be used in the next step without purification. [ES-MS] (ESI+) calculated for $C_9H_{10}N_2OS$ $[M+H]^+$ 195.05; found 195.0.

**2,2-dibromo-1-(2,6-dichlorophenyl)ethanone (2)**

To a solution of 1-(2,6-dichlorophenyl)ethanone (1.275 g, 6.74 mmol) in anhydrous THF (6.74 mL), was added portionwise phenyltrimethylammonium tribromide (5.068 g, 13.48 mmol) over a period of one hour. The solution was stirred under argon overnight. Cold water (5 mL) was added and the solution was extracted with dichloromethane. The combined organic layers were washed with brine, dried over magnesium sulfate, and concentrated. The crude solid was purified by flash chromatography (silica gel, hexanes/ethyl acetate) to afford 2,2-dibromo-1-(2,6-dichlorophenyl)ethanone **2** as a clear gel (2.363 g, quant.). [ES-MS] (ESI+) calculated for $C_8H_4Br_2Cl_2O$ $[M-H]^-$ 344.8; found 344.6, 346.6.

**5-(2,6-dichlorophenyl)-1,2,4-triazin-3-amine (3)**

Under argon, morpholine (2.46 mL, 28.47 mmol) was added all at once to a solution of 2,2-dibromo-1-(2,6-dichlorophenyl)ethanone (2.363 g, 6.81 mmol) in anhydrous THF (2.27 mL). The resulting mixture was thoroughly degassed and then heated slowly over one hour to 67 °C. The solution was kept stirring at this temperature for 2 days after which it was cooled to room temperature. The white solid was filtered off and the residue was concentrated. Under argon, the crude residue was then dissolved in methanol (11.16 mL), aminoguanidine bicarbonate (0.955 g, 7.01 mmol) was added followed by acetic acid (1.17 mL). The resulting mixture was briefly degassed and stirred at room temperature for 1.5 hours, then heated at 67 °C for 2 days. The reaction mixture was then cooled to room temperature, concentrated, and purified by flash chromatography (silica gel, dichloromethane/methanol) to afford 5-(2,6- dimethoxyphenyl)-1,2,4-

triazin-3-amine **3** as a brown oil (0.246 g, 15%). [ES-MS] (ESI+) calculated for $C_9H_6Cl_2N_4$ $[M+H]^+$ 241.00; found 240.9, 242.9.

**3-chloro-5-(2,6-dichlorophenyl)-1,2,4-triazine (4)**

Under argon, a solution of tert-butyl nitrite (0.68 mL, 5.72 mmol) and copper(II) chloride (0.563 g, 4.19 mmol) in anhydrous acetonitrile (1 mL) was heated at 65 °C, and a solution of 5-(2,6- dimethoxyphenyl)-1,2,4-triazin-3-amine **3** (0.919 g, 3.81 mmol) in anhydrous acetonitrile (14.66 mL) and dichloromethane (9.77 mL) was added. The reaction mixture was stirred at 65 °C for 1.5 hours. The solution was then poured into aqueous hydrochloric acid 6N cooled to 0 °C. The mixture was extracted with diethyl ether, combined organic extracts were washed with brine, then dried over magnesium sulfate, filtered, and evaporated. The residue was purified by flash chromatography (silica gel, hexane/ethyl acetate) to afford 3-chloro-5-(2,6-dichlorophenyl)-1,2,4-triazine **4** as a yellow solid (0.482 g, 49%). [ES-MS] (ESI+) calculated for $C_9H_6Cl_3N_3$ $[M+H]^+$ 259.95; found 259.9, 261.9.

**5-(2,6-dichlorophenyl)-*N*-((5-methoxybenzo[*d*]thiazol-2-yl)methyl)-1,2,4-triazin-3-amine (5)**

Under argon, (5-methoxy-1,3-benzothiazol-2-yl)methylamine (0.395 g, 2.04 mmol) and DIPEA (0.35 mL) were added to a solution of 3-chloro-5-(2,6-dichlorophenyl)-1,2,4-triazine (0.482 g, 1.85 mmol) in anhydrous acetonitrile (5.61 mL), and the reaction mixture was refluxed overnight. Water was added and the solution was extracted with ethyl acetate. Combined organic extracts were washed with water, dried over magnesium sulfate, filtered, and concentrated. The residue was purified by flash chromatography (silica gel, hexane/ethyl acetate) to give 5-(2,6-dichlorophenyl)-*N*-((5-methoxy-1,3-benzothiazol-2-yl)methyl)-1,2,4-triazin-3-amine **5** as a beige solid (0.603 g, 78%). [ES-MS] (ESI+) calculated for $C_{18}H_{13}Cl_2N_5OS$ $[M+H]^+$ 418.03; found 418.0, 419.0.

**2-(((5-(2,6-dichlorophenyl)-1,2,4-triazin-3-yl)amino)methyl)-1,3-benzothiazol-5-ol (6)**

Under argon, a solution of boron tribromide in dichloromethane (1M, 13.8 mL, 13.8 mmol) was slowly added to a solution of 5-(2,6-dichlorophenyl-*N*-((5-methoxy-1,3-benzothiazol-2-yl)methyl)-1,2,4-triazin-3-amine (0.576 g, 1.38 mmol) in anhydrous dichloromethane (13.8 mL) at -78 °C. The reaction mixture was stirred overnight allowing the temperature to rise to room temperature. Then, the solution was cooled to 0 °C and methanol (4.6 mL) was added. The resulting mixture was stirred at 50 °C for 30 minutes then concentrated. Ethyl acetate and water were added to the residue. The solution was extracted with ethyl acetate; combined organic extracts were dried over magnesium sulfate, filtered, and evaporated. The crude residue was purified by flash chromatography (silica gel, dichloromethane/methanol) to afford 2-(((5-(2,6-dichlorophenyl)-1,2,4-triazin-3-yl)amino)methyl)-1,3-benzothiazol-5-ol **6** as a yellow solid (0.528 g, 95%). [ES-MS] (ESI+) calculated for $C_{17}H_{11}Cl_2N_5OS$ $[M+H]^+$ 404.1; found 403.9, 405.9.

**ethyl 2-((2-(((5-(2,6-dichlorophenyl)-1,2,4-triazin-3-yl)amino)methyl)-1,3-benzothiazol-5-yl)oxy)acetate (7)**

Under argon, a solution of 2-(((5-(2,6-dichlorophenyl)-1,2,4-triazin-3-yl)amino)methyl)-1,3-benzothiazol-5-ol (0.528 g, 1.31 mmol) and potassium carbonate (0.362 g, 2.62 mmol) in acetone (10.92 mL) was stirred at room temperature for 30 minutes. Ethyl bromoacetate (0.29 mL, 2.62 mmol) was added and the reaction mixture was stirred at 60°C overnight. The mixture was filtered, rinsed with methanol and dichloromethane, and the filtrate was evaporated. The crude residue was purified by flash chromatography (silica gel, dichloromethane/methanol) to afford ethyl 2-((2-(((5-(2,6-dichlorophenyl)-1,2,4-triazin-3-yl)amino)methyl)-1,3-benzothiazol-5-yl)oxy)acetate **7** as a yellow solid (0.370 g, 58%). [ES-MS] (ESI+) calculated for $C_{21}H_{17}Cl_2N_5O_3S$ $[M+H]^+$ 490.05; found 490.0, 492.0.

**2-((2-(((5-(2,6-dichlorophenyl)-1,2,4-triazin-3-yl)amino)methyl)-1,3-benzothiazol-5-yl)oxy)acetic acid (8)**

Lithium hydroxide (0.053 g, 2.23 mmol) was added to a solution of ethyl 2-((2-(((5-(2,6-dichlorophenyl)-1,2,4-triazin-3-yl)amino)methyl)-1,3-benzothiazol-5-yl)oxy)acetate (0.370 g, 0.75 mmol) in THF (9.32 mL) and water (4.66 mL). The reaction was stirred vigorously under argon at room temperature for 1.5 hours. Hydrochloric acid 1N was added and the solution was extracted with diethyl ether and ethyl acetate. The combined organic extracts were dried over magnesium sulfate, filtered and evaporated. The crude residue was purified by flash chromatography (silica gel, dichloromethane/methanol) to afford 2-((2-(((5-(2,6-dichlorophenyl)-1,2,4-triazin-3-yl)amino)methyl)-1,3-benzothiazol-5-yl)oxy)acetic acid **8** as a beige solid (0.297 g, 85%). [ES-MS] (ESI+) calculated for $C_{19}H_{13}Cl_2N_5O_3S$ $[M+H]^+$ 462.02, found 461.9, 464.0.

**2-((2-(((5-(2,6-dichlorophenyl)-1,2,4-triazin-3-yl)amino)methyl)-1,3-benzothiazol-5-yl)oxy)-*N*-(2-hydroxyethyl)acetamide (9)**

Under argon, ethanolamine (4.70 µL, 0.079 mmol) and DIPEA (33.57 µL) were added to a solution of 2-((2-(((5-(2,6-dichlorophenyl)-1,2,4-triazin-3-yl)amino)methyl)-1,3-benzothiazol-5-yl)oxy)acetic acid (30 mg, 0.064 mmol) and HATU (37.90 mg, 0.1 mmol) in anhydrous dichloromethane (0.71 mL). The reaction mixture was stirred for 1 hour at room temperature. Water was added and the solution was extracted with ethyl acetate. The combined organic extracts were washed with brine, dried over magnesium sulfate, filtered, and evaporated. The crude solid was purified by preparative HPLC (0-40% acetonitrile in water + 0.1% TFA) to afford 2-((2-(((5-(2,6-dichlorophenyl)-1,2,4-triazin-3-yl)amino)methyl)-1,3-benzothiazol-5-yl)oxy)-*N*-(2-hydroxyethyl)acetamide **9** as a white solid (10 mg, 31%). Purity was confirmed by analytical HPLC (Agilent Poroshell 120 EC-C18 column, 2.7 µm, 3.0 x 50 mm; water + 0.1% formic acid/acetonitrile + 0.1% formic acid; 0-100% gradient over 10 min; flow rate 0.5 mL/min). HRMS (ESI+) calculated for $C_{21}H_{18}Cl_2N_6O_3S$ $[M+H]^+$ 505.0611; found 505.0611, 507.0585.

**HPLC purity of 1.**

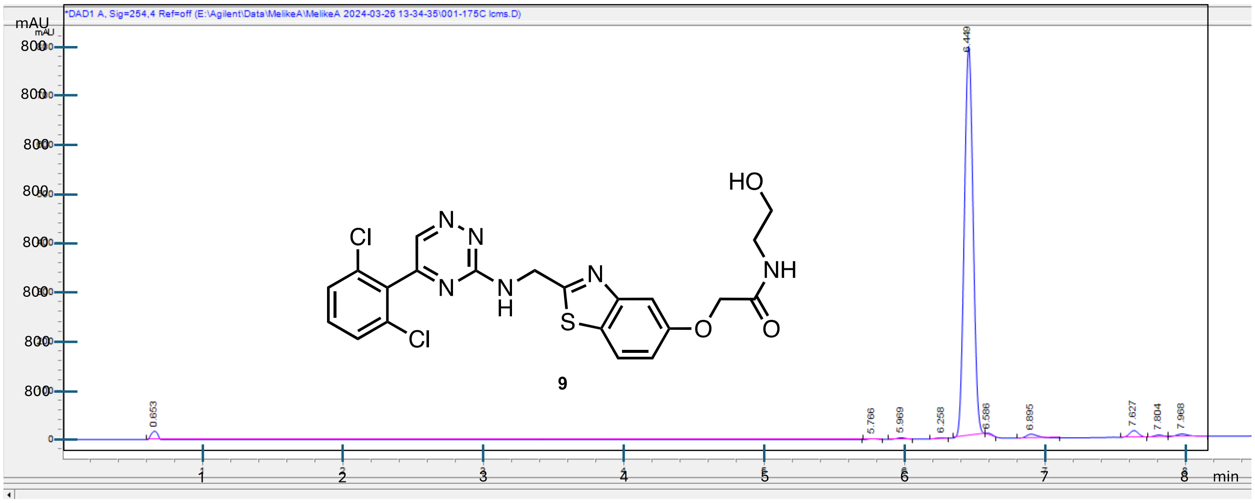

**Synthesis of LPC-058**.

The synthesis of LPC-058 was adapted from Liang et al (5).

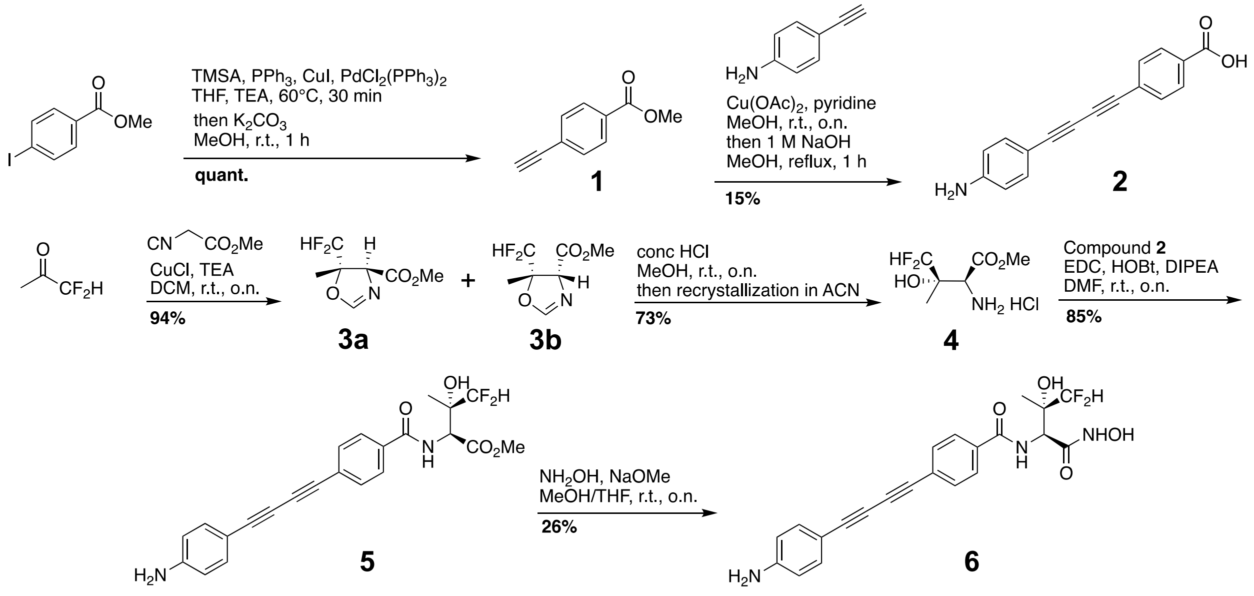


**Scheme S3: Synthetic scheme for the LpxC inhibitor LPC-058.**

**4-Ethynylbenzoic acid methyl ester (1)**

4-iodobenzoic acid methyl ester (500 mg, 1.91 mmol), $PPh_3$ (50 mg, 0.191 mmol), CuI (36 mg, 0.191 mmol), and $PdCl_2(PPh_3)_2$ (67 mg, 0.095 mmol) were combined under an inert atmosphere. Anhydrous THF (16 mL) was added followed by TEA (16 mL) and TMSA (0.26 mL, 1.91 mmol), and the reaction mixture was stirred at 60°C for 30 mins. The reaction was then quenched with 10% $NH_4Cl$ and extracted three times with ethyl acetate. The combined organic layers were washed with brine and dried over $MgSO_4$. The solvent was removed under reduced pressure, the resulting residue was dissolved in MeOH (12 mL), and $K_2CO_3$ (1.32 g, 9.57 mmol) was added. The reaction was stirred until the deprotection was complete as determined by TLC. The solution was filtered, and the solvent was removed under reduced pressure to afford the desired product, which was used without further purification. Yield: quant. $^1$H NMR (400 MHz, $CDCl_3$) δ 8.01 (d, $J$ = 8.4 Hz, 2H), 7.57 (d, $J$ = 8.4 Hz, 2H), 3.94 (s, 3H), 3.25 (s, 1H).

**4-((4-Aminophenyl)buta-1,3-diyn-1-yl)benzoic acid (2)**

Compound **1** (86 mg, 0.54 mmol), 4-ethynylaniline (313 mg, 2.67 mmol), and $Cu(OAc)_2$ (196 mg, 1.08 mmol) were combined under an inert atmosphere. Pyridine (2.1 mL) and MeOH (2.1 mL) were added, and the reaction was stirred at room temperature overnight. Water was then added, and the mixture was extracted with ethyl acetate. The organic layer was washed with brine, dried with $MgSO_4$, and the solvent was removed under reduced pressure. The residue was then redissolved in 1 M NaOH and heated to reflux for 1 h. The reaction was cooled to room temperature, and the pH of the solution was adjusted to 3 using concentrated HCl. The resulting precipitate was collected and used without further purification. Yield: 15% [ES-MS] (ESI+) calculated for $C_{17}H_{11}NO_2$ $[M + H]^+$ 262.3; found 262.0.

**Methyl (4*S*/*R*,5*S*)-5-(difluoromethyl)-5-methyl-4,5-dihydrooxazole-4-carboxylate (3a/3b)**

CuCl (10 mg, 0.10 mmol) was sealed with a stir bar under an inert atmosphere. 1,1-Difluoroacetone (0.20 mL, 3.0 mmol), TEA (0.015 mL, 0.125 mmol), and DCM (3.5 mL) were added, and the reaction mixture was cooled to 0°C. Methyl 2-isocyanoacetate (0.25 mL, 2.5 mmol) was added dropwise and stirred overnight at room temperature. The reaction was diluted with DCM and washed with a 10% ammonia solution three times. The organic layer was washed with brine and dried with $MgSO_4$. The solvent was removed under reduced pressure, and the resulting mixture of diastereomers were used without further purification. Yield: 94% [ES-MS] (ESI+) calculated for $C_7H_9F_2NO_3$ $[M + H]^+$ 194.2; found 194.0.

**Methyl (2*S*,3*S*)-2-amino-4,4-difluoro-3-hydroxy-3-methylbutanoate hydrochloride (4)**

A mixture of diastereomers **3a** and **3b** (97 mg, 0.50 mmol) were dissolved in MeOH (0.50 mL) under an inert atmosphere, and the reaction was placed in a water bath to maintain the temperature as concentrated HCl (0.10 mL) was added dropwise. The reaction mixture was stirred at room temperature overnight, and then the solvent was removed under reduced pressure. The resulting residue was diluted with MTBE and stirred vigorously for 2 hours. The MTBE was removed under reduced pressure, and the resulting solid was recrystallized in ACN. The crystals were collected and used without further purification. Yield: 73% $^1$H NMR (400 MHz, MeOD) δ 6.00 (t, *J* = 55.1 Hz, 1H), 4.20 (s, 1H), 3.90 (s, 3H), 1.34 (t, *J* = 1.7 Hz, 3H).

**Methyl (2S,3S)-2-(4-((4-aminophenyl)buta-1,3-diyn-1-yl)benzamido)-4,4-difluoro-3-hydroxy-3-methylbutanoate (5)**

Compound **2** (21 mg, 0.08 mmol), compound **4** (15 mg, 0.084 mmol), EDC•HCl (18 mg, 0.096 mmol), and HOBt hydrate (13 mg, 0.096 mmol) were dissolved in DMF (0.84 mL), and the reaction mixture was cooled to 0°C. DIPEA (0.06 mL, 0.32 mmol) was added, and the reaction was stirred at room temperature overnight. The reaction was then quenched with water and extracted with ethyl acetate three times. The combined organic layers were washed with brine and dried with $MgSO_4$. The solvent was removed under reduced pressure, and the resulting solid was purified by column chromatography ($SiO_2$; ethyl acetate/hexanes; 0-100% gradient) Yield: 85% [ES-MS] (ESI+) calculated for $C_{23}H_{20}F_2N_2O_4$ $[M + H]^+$ 427.4; found 427.1.

**4-((4-Aminophenyl)buta-1,3-diyn-1-yl)-*N*-((2*S*,3*S*)-4,4-difluoro-3-hydroxy-1-(hydroxyamino)-3-methyl-1-oxobutan-2-yl)benzamide (6)**

Compound **5** (29 mg, 0.068 mmol) was dissolved in MeOH (0.11 mL) and THF (0.11 mL) under an inert atmosphere and put on ice. $NH_2OH$•HCl (24 mg, 0.34 mmol) was added followed by a 0.5 M NaOMe solution in MeOH (1.0 mL, 0.51 mmol). The reaction was stirred at room temperature overnight and was then quenched with saturated $NH_4Cl$ and extracted with ethyl acetate three times. The combined organic layers were washed with brine and dried with $MgSO_4$. The solvent was removed under reduced pressure, and the resulting residue was purified by preparative HPLC using a Shimadzu LC20 Series preparative HPLC, Phenomenex Kinetex C18 column, 5 µm, 250 x 21.2 mm; water + 0.1% TFA/acetonitrile + 0.1% TFA; 0-100% gradient over 30 min; flow rate 15 mL/min. The product peak was collected at 10 min and lyophilized to produce the desired product as a mixture of rotamers in the form of a brown oil. Purity was confirmed by analytical HPLC (Agilent Poroshell 120 EC-C18 column, 2.7 µm, 3.0 x 50 mm; water + 0.1% formic acid/acetonitrile + 0.1% formic acid; 0-100% gradient over 10 min; flow rate 0.5 mL/min). Yield: 26% [ES-MS] (ESI+) calculated for $C_{22}H_{20}F_2N_3O_4$ $[M + H]^-$ 428.1416; found 428.1420. $^1H$ NMR (400 MHz, MeOD, mixture of rotamers) δ 7.90 (d, $J$ = 8.4 Hz, 1H), 7.71 (d, $J$ = 8.5 Hz, 2H), 7.67 (d, $J$ = 8.4 Hz, 2H), 7.40 (d, $J$ = 8.6 Hz, 1H), 5.85 (t, $J$ = 56.0, 0.5H), 5.82 (t, $J$ = 56.0, 0.5H), 4.81 (s, 0.5H), 4.75 (s, 0.5H), 1.38 (s, 1.5H), 1.35 (s, 1.5H).

**HPLC purity of LPC-058.**

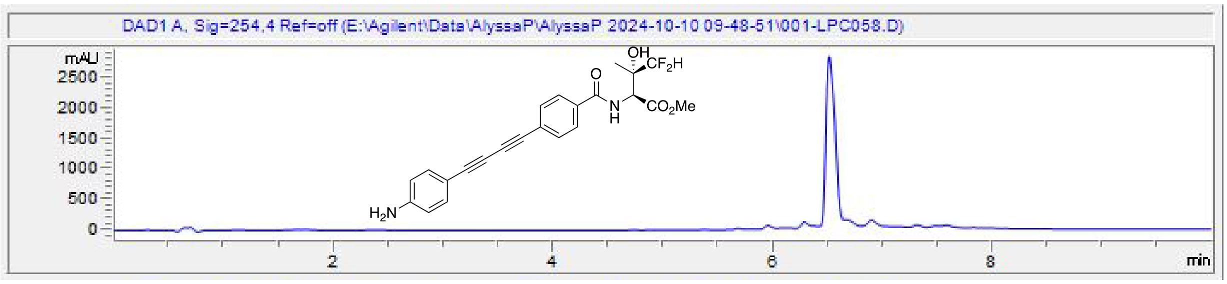
